# Genetic changes linked to two different syndromic forms of autism enhance reinforcement learning in early adolescent male but not female mice

**DOI:** 10.1101/2025.01.15.633099

**Authors:** Juliana Chase, Jing-Jing Li, Wan Chen Lin, Lung-Hao Tai, Fernanda Castro, Anne GE Collins, Linda Wilbrecht

## Abstract

Autism Spectrum Disorder (ASD) is characterized by restricted and repetitive behaviors and social differences, both of which may manifest, in part, from underlying differences in corticostriatal circuits and reinforcement learning. Here, we investigated reinforcement learning in developing mice with mutations in either *Tsc2* or *Shank3*, both high-confidence ASD risk genes associated with major syndromic forms of ASD. Using an odor-based two-alternative forced choice (2AFC) task, we tested early adolescent mice of both sexes and found male *Tsc2* and *Shank3B* heterozygote (Het) mice showed enhanced learning performance compared to their wild type (WT) siblings. No gain of function was observed in females. Using a novel reinforcement learning (RL) based computational model to infer learning rate as well as policy-level task engagement and disengagement, we found that the gain of function in males was driven by an enhanced positive learning rate in both *Tsc2* and *Shank3B* Het mice. The gain of function in Het males was absent when mice were trained with a probabilistic reward schedule. These findings in two ASD mouse models reveal a convergent learning phenotype that shows similar sensitivity to sex and environmental uncertainty. These data can inform our understanding of both strengths and challenges associated with autism, while providing further evidence that sex and experience of uncertainty modulate autism-related phenotypes.

**Significance Statement:** Reinforcement learning is a foundational form of learning that is widely used in behavioral interventions for autism. Here, we measured reinforcement learning in developing mice carrying genetic mutations linked to two different syndromic forms of autism. We found that males showed strengths in reinforcement learning compared to their wild type siblings, while females showed no differences. This gain of function in males was no longer observed when uncertainty was introduced into the reward schedule for correct choices. These findings support a model in which diverse genetic changes interact with sex to generate common phenotypes underlying autism. Our data further support the idea that autism risk genes may produce strengths as well as challenges in behavioral function.

## Introduction

Autism Spectrum Disorder (ASD) is a common neurodevelopmental condition that is characterized by restricted and repetitive behaviors in addition to differences in social communication. Autism is genetically heterogeneous, but it is thought that mutations to single genes can cause disruptions in brain development, leading to the condition. These genetic differences can be introduced into mouse lines for study (Tsai et al., 2012; Peça et al., 2011).

Restricted and repetitive behaviors observed in autism are likely related to differences in learning and corticostriatal circuit function (Fuccillo, 2016). Differences in these circuits may also influence social learning (Matthiesen et al., 2023). Discrete regions of the striatum are thought to support both cognitive and motor learning with separate cortical, limbic, and neuromodulatory inputs controlling different aspects of action generation, cue and action evaluation, and reinforcement (Cox & Witten, 2019; Peters et al., 2021). Underscoring the striatum’s critical role in ASD, studies consistently link changes in striatal development and function with ASD symptoms and severity (Qiu et al., 2016; Fuccillo, 2016; Rothwell, 2016; Comparan-Meza et al., 2021; Dölen & Malenka, 2014). Mutations to autism risk genes have also been shown to alter corticostriatal circuit function and structure (Rothwell et al., 2014; Platt et al., 2017; Le Merrer et al., 2023; Cording & Bateup, 2023). Notably, many mouse models with mutations to ASD risk genes show enhanced motor learning on the rotarod task, a well-established assay of striatal function (Cording & Bateup, 2023). However, whether these gains extend to reinforcement learning tasks, which also depends on the striatum (Peters et al., 2021; Reinhold et al., 2023), remains unclear.

Here we focus on learning in two lines of mice, *Tsc2^+/-^*and *Shank3B*^+/-^, that have been engineered to mimic genetic differences found in two syndromic forms of autism: Tuberous Sclerosis Complex (TSC) and Phelan-McDermid Syndrome, respectively. Syndromic forms of autism, including TSC and Phelan-McDermid Syndrome, account for around 1% of *de novo* genetic causes of ASD (de la Torre-Ubieta et al., 2016). TSC is defined by a mutation to either the *TSC1* or *TSC2* gene. People with either TSC mutation often exhibit brain lesions called tubers, epilepsy, and cognitive deficits of varying degrees, and, in 25-60% of cases, they also receive an ASD diagnosis (Gipson et al., 2019; Boronat, Thiele, & Caruso, 2017; Baker et al., 1998; Capal et al., 2021). The protein products of the *TSC1* and *TSC2* genes, hamartin and tuberin respectively, form a protein complex that negatively regulates the mammalian target of rapamycin complex 1 (mTORC1) (Huang & Manning, 2008). mTORC1 is a central and conserved intracellular signaling pathway that coordinates essential cellular processes such as autophagy, cellular metabolism, and protein and lipid synthesis (Laplante & Sabatini, 2012). Phelan-McDermid Syndrome is driven by either a deletion of the terminal end of chromosome 22 or a mutation to *SHANK3* (Costales & Kolevzon, 2015). It is characterized by intellectual disability, neuromuscular effects, and is accompanied by an autism diagnosis in approximately 85% of cases (Durand et al., 2006) which account for up to 2% of all ASD cases (Leblond et al., 2014). The protein encoded by *SHANK3,* SH3 and multiple ankyrin repeat domains 3 (SHANK3), is one of a family of proteins that bind directly with synapse-associated protein 90/postsynaptic density 95-associated protein (SAPAP) and postsynaptic density protein (PSD) 95 (Naisbitt et al., 1999; Kim & Sheng, 2004). Together they form the PSD-95-SAPAP-SHANK complex and play an integral role in synaptic formation and development (Sala et al., 2001; Baron et al., 2006; Peça et al., 2011). SHANK3 is predominantly found in the striatum and is localized postsynaptically at glutamatergic synapses onto spiny projection neurons (SPNs) (Peça et al., 2011), synapses known to be important for learning (Hwang et al., 2022). Mice with a deletion of the PDZ region of Shank3, called *Shank3B^-/-^*, conditionally knocked out of indirect pathway SPNs (iSPNs) show increased grooming, a repetitive behavior that can be readily assessed in rodents (Wang et al., 2017). Importantly, haploinsufficiency of *SHANK3* in Phelan-McDermid patients has been shown to be responsible for the neurological symptoms associated with the syndrome (Durand et al., 2006).

TSC1/2 and SHANK3 have distinct cellular functions, yet they may have a convergent impact on the development of circuits, including corticostriatal circuits which are implicated in cognitive and motor learning (Thorn et al., 2010; Koralek et al., 2012; Fisher et al., 2020; Peters et al., 2021; Reinhold et al., 2023; Cording and Bateup, 2023). Importantly, *Tsc1* and *Shank3* manipulations in mice have both been shown to enhance cortical connectivity onto striatal spiny projection neurons (SPNs) (Benthall et al., 2018; Peixoto et al., 2016;2019; Benthall et al., 2021). In *Shank3B^-/-^* mice, hyperconnectivity was greater earlier in development and transitioned to hypoconnectivity in adulthood (Peixoto et al., 2019). This convergent circuit phenotype could plausibly lead to convergent behavioral differences, particularly in diverse forms of learning supported by striatal functions. Therefore, we chose to examine reinforcement learning in developing *Tsc2* and *Shank3B* mice during the first days of acquiring correct action-odor cue associations.

Our goal was to test whether developing mice, haploinsufficient for *Tsc2* or *Shank3B,* would show a gain or loss of function in an odor-based two alternative forced choice (2AFC) task where they were trained to associate an odor with a correct action to receive a water reward. Based on previous rotarod data, we may have expected *Tsc2* Het animals to show strong motor coordination, but whether reinforcement learning is stronger or weaker than WT was unknown.

We chose to focus on initial stages of learning in developing mice (starting odor learning around postnatal day (P)30) to enhance translational value. In humans, development is a prime period for intervention and the numbers of trials for training are more limited. Yet, in animal studies, rodents with altered autism risk genes are typically tested in adulthood and examined after extensive overtraining.

We also chose to study both male and female mice and to separately analyze results in each sex. Gene expression unfolds within a larger context of the developing body, affecting cellular and behavioral phenotypes in dynamic ways, particularly in adolescence when puberty is beginning (Ferri et al., 2018; Delevich et al., 2021; Larsen and Luna, 2018). There has historically been a male bias in autism diagnoses (Baio et al., 2018), and male biased sex differences are also observed in mice with mutations to ASD risk genes (Ferri et al., 2018; Grissom et al., 2018). However, in the past few decades, the male:female ratio is decreasing in human diagnosis (Hull, Petrides, & Mandy, 2020) and there is growing awareness of a need to understand putative female resilience and/or how ASD phenotypes may present differently in females.

To examine the latent processes that mice use to solve the odor-based 2AFC task, we fit trial-by-trial data using multiple reinforcement learning (RL) models, including a novel hybrid model that we recently developed that accounts for learning and exploration noise, as well as periods of disengagement at the policy level (Li et al., 2024).

## Results

### Early adolescent male *Tsc2* Het mice show gain of function in early learning

We first tested early adolescent male mice with a loss-of-function mutation in *Tsc2* in the odor-based 2AFC. To begin, *Tsc2* Het (n = 15) and WT (n = 13) mice were water restricted and habituated to the chamber, the central initiation port, and its two peripheral water delivery ports. After 4 days of habituation (Figure 1A), the mice were exposed to two odors (odors “A” and “B”) that were stably paired with a water reward on either the left or right peripheral port (Figure 1B). In the first odor learning session (Session 1), male *Tsc2* Hets steadily increased performance throughout the session to reach over 70% correct, on average, in the last quarter of the session, whereas their WT littermates maintained performance near to chance (Figure 1C; 2-way RM ANOVA: geno: *F*(1, 26) = 7.27, *p* = 0.01; time: *F*(1.75, 45.55) = 15.05, *p* < 0.0001; geno x time: *F*(3, 78) = 3.92, *p* = 0.01; Tukey post-hoc test in quartile 4 showed Hets significantly higher than WTs (*p* < 0.01)). In the second session of odor learning (the following day), Het animals again significantly outperformed their WT littermates by choosing the correct side more consistently following each odor presentation (Figure 1D; 2-way RM ANOVA: geno: *F*(1, 26) = 8.56, *p* = 0.007; time: *F*(2.165, 56.29) = 10.05, *p* = 0.0001; geno x time: *F*(3, 78) = 1.17, *p* =0.16; significant post-hoc differences in quartiles 2, 3, and 4).

**Figure 1:**
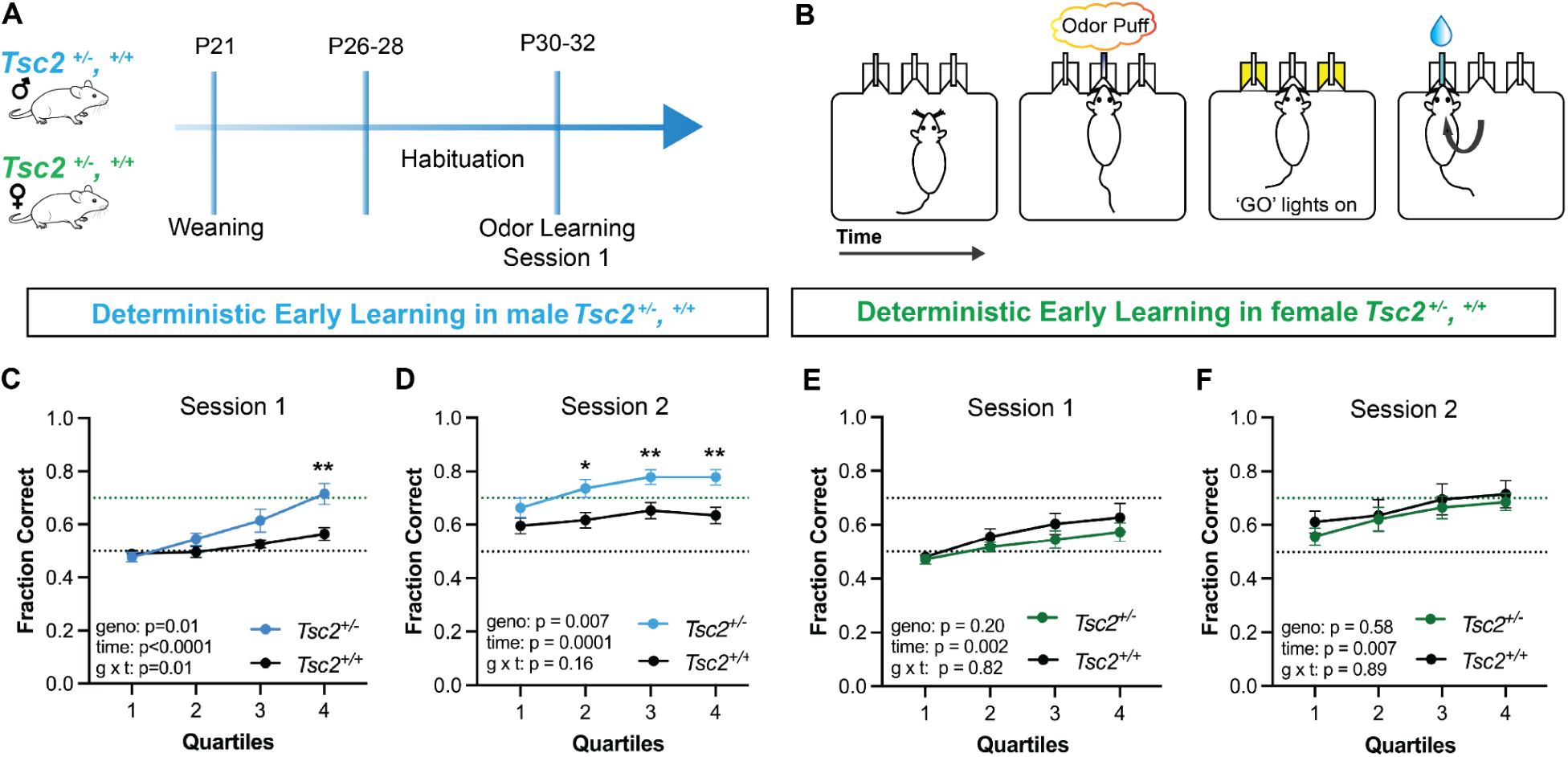
Early adolescent male, but not female *Tsc2* Het mice show a gain of function in learning in early training in an odor based 2AFC task. (A) Timeline of training. One week after weaning, male and female Tsc2 Het and WT animals were habituated to the chamber and ports through a shaping procedure in which they first learned to poke into the center port, followed by the peripheral ports, to receive water rewards. Here we used a deterministic reward schedule, where every correct choice led to a water reward. Odor cues were not present during habituation. At P30-P32 mice began session 1 of the odor-based 2AFC task. (B) In this task, mice self-initiate a trial by nose poke into the center port, which releases one of two odor cues. Bilateral “Go” lights then indicate the availability of reward, prompting mice to choose a peripheral side port. Choosing the correct port for that odor, results in a 2µL water reward. (C-D) Performance in male *Tsc2* Het mice was significantly greater than WT males in both the first and second session of odor learning. This was captured by a significant main effect of genotype in both sessions and a genotype x time interaction in session 1. Significant Tukey post-hoc tests between Het and WT for each quartile are indicated by asterisks. (E-F) We also tested female *Tsc2* Het and WT mice in the same odor-based 2AFC task on the same schedule. Female *Tsc2* Het and WT did not show differences in learning performance for Session 1 or Session 2. Full statistical results are reported in the main text. Male *Tsc2* Het, n = 15; WT, n = 13; Female *Tsc2* Het, n = 10; WT, n = 10. Learning curves show mean for each quartile of all trials performed by individual mice ± SEM. \**p*<0.05, \*\**p*<0.01.

In addition to measuring performance, we quantified the trial reinitiation times using the intertrial interval (ITI) – the time from the reward port to the center port – as a reaction-time-like metric that may reflect differences in motivation, motor coordination, or both. In session 1, the ITI metric was comparable between WT and Het, but in session 2 Het mice were faster than WT (Supplemental Figure 1F: Unpaired *t*-test: Session 1: *t*(26) = 1.29, *p* = 0.20; Session 2: *t*(25) *=* 2.92, *p* = 0.007). We observed a significant negative correlation between performance and ITI in quartiles in both *Tsc2* Het and WT groups in the second session (Spearman’s correlation: WT: *r* = −0.52, *p* < 0.0001; HET: *r* = −0.42, *p* = 0.0004). Together, these data reveal a gain of function in *Tsc2* males in the earliest sessions of odor-action reinforcement learning.

### Early adolescent female *Tsc2* Het mice have no gain of function in early learning

To test if there was a similar or different phenotype in *Tsc2* females, we trained female *Tsc2* Het (n = 10) and WT (n = 10) mice in the same odor 2AFC task. *Tsc2* Het and WT females learned the odor-based 2AFC task at a comparable rate (Figure 1E: Session 1: 2-way RM ANOVA: geno: *F*(1, 18) = 1.69, *p* = 0.20; time: *F*(1.90, 34.31) = 7.5, *p* = 0.002; geno x time: *F*(3, 54) = 0.30, *p* =0.82) and both *Tsc2* Het and *Tsc2* WT groups reached 70% average performance by the end of the second session (Figure 1F, Session 2: 2-way RM ANOVA: geno: *F*(1, 18) = 0.31, *p* = 58; time: *F*(2.18, 39.33) = 8.42, *p* = 0.0007; geno x time: *F*(3, 54) = 0.25, *p* = 0.89).

Trial reinitiation times, quantified by ITI, were also comparable between female Tsc2 Het and WT animals across both sessions (Supplemental Figure 1G: Session 1, Mann-Whitney test: *U* = 41, *p* = 0.52; Session 2, Unpaired *t*-test: *t*(16) = 0.22, *p* = 0.82). Performance in session 2 was also negatively correlated with ITI in both genotypes (Pearson’s correlation: WT: *r* = –0.91, *p* < 0.0001; HET: *r* = –0.47, *p* = 0.002).

To put these sex differences in context we also compared males and females in a supplemental analysis. *Tsc2* WT males and WT females had comparable performance in both sessions (Supplemental Figure 1B-C, 2-way RM ANOVA: Session 1: sex: *F*(1, 21) = 3.21, *p* = 0.08; time: *F*(1.70, 35.75) = 8.18, *p* = 0.001; sex x time: *F*(3, 63) = 1.337, *p* = 0.27; Session 2: sex: *F*(1, 21) = 0.58, *p* = 0.45; time: *F*(2.40, 50.52) = 5.86, *p* = 0.003; sex x time: *F*(3, 63) = 1.01, *p* = 0.39), while female *Tsc2* Het and male *Tsc2* Het showed different learning across both sessions with Het males showing a significant gain of function over Het females (Supplemental Figure 1D-E, 2-way RM ANOVA: Session 1: sex: *F*(1, 23) = 4.33, *p* =0.04; time: *F*(1.79, 41.39) = 12.74, *p* < 0.0001; sex x time: *F*(3, 69) = 2.16, *p* =0.10; Session 2: sex: *F*(1, 23) = 6.17, *p* = 0.02; time: *F*(2, 46.11) = 13.77, *p* < 0.0001; sex x time: *F*(3, 69) = 0.11, *p* = 0.95). As a control, we also tested whether daily weight of male and female *Tsc2* Het and WT mice was correlated with performance. We found no evidence of a significant relationship between these variables in either sex (Supplemental Figure 1H-K).

### Gain of function in developing*Tsc2* Het males is captured by higher learning rates

We next turned to computational modeling to see whether an exploration of latent variables in trial-by-trial learning could help inform which cognitive mechanisms were responsible for differences in performance between male *Tsc2* Het and WT mice. This also allowed us to explore potential latent differences in females. We compared five models that extended a basic reinforcement learning (RL) framework that posits, based on research from our lab (Chase et al., 2024), that mice rely on RL in an odor-based associative learning task. RL assumes that animals use reward outcomes to estimate the value of making a choice and iteratively update the values by the discrepancy between an obtained and an expected reward. To test if performance differences emerged from differences in attention or task disengagement, we also investigated a hybrid model we recently developed (Li et al., 2024). In this model, the agent alternates between two latent states that govern distinct policies: an RL-based policy when in an “engaged” state and a non-RL, side-biased policy when in a “disengaged” latent state (probabilistically choosing a single side while disregarding trial stimuli and outcomes). A hidden Markov model regulates transitions between these states (Figure 2A).

**Figure 2:**
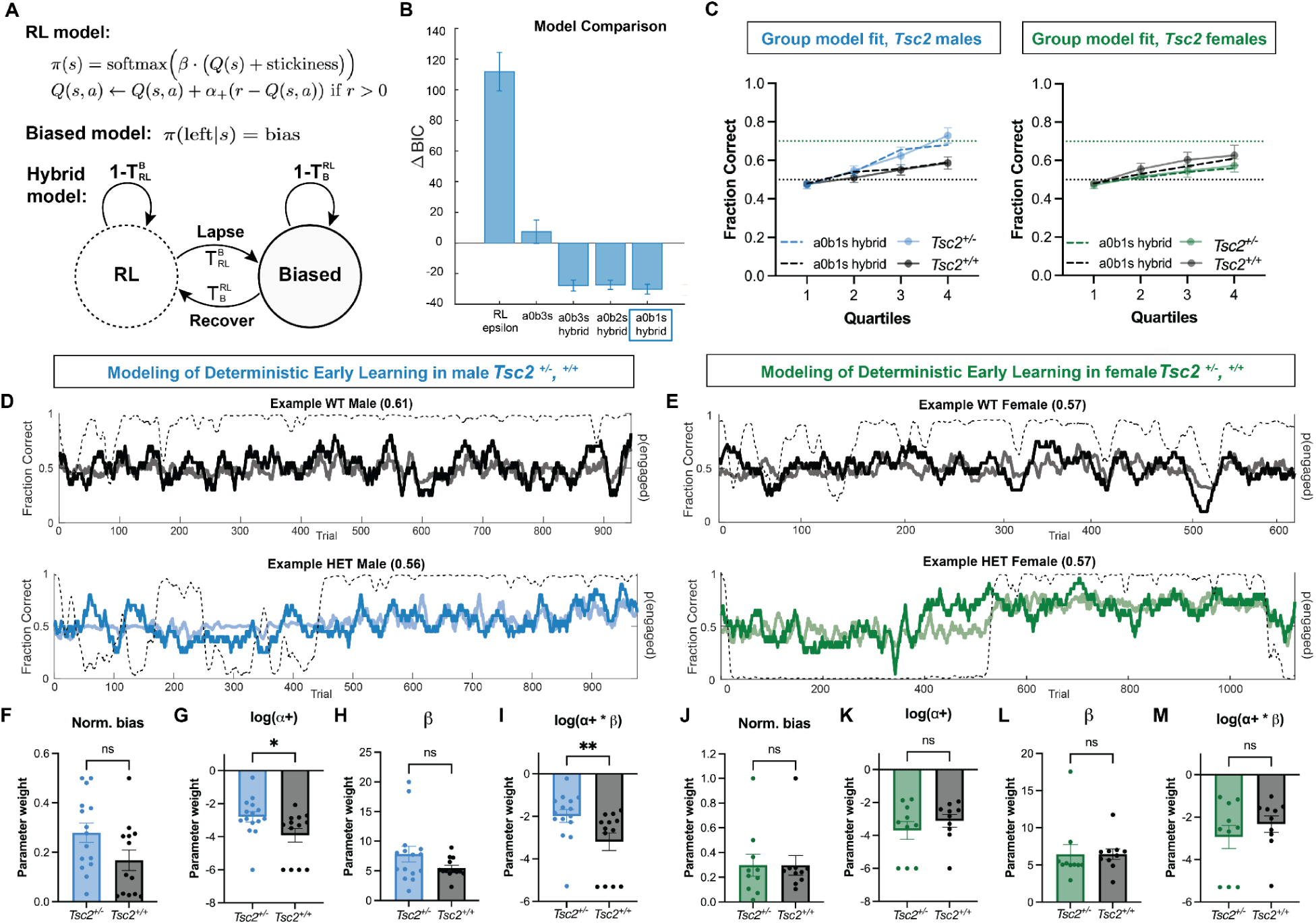
Early adolescent male, but not female, *Tsc2* Het mice have a higher learning rate when fit with the winning hybrid model. We examined a series of models that use a reinforcement learning (RL) framework to account for differences in choice behavior between genotypes in Session 1 (see Materials & Methods). (A) Several of the candidate models that were tested were mixture models that combined a standard Q-learning RL model, representing an engaged state, with a biased model, representing a disengaged state, to account for periods of task disengagement at the policy level. The transition probabilities of these separate policy states are governed by a hidden Markov model. (B) Using Bayesian Information Criterion (BIC), we determined that models with the mixture policy (named hybrid here) had a lower BIC score. The change in BIC is calculated by the difference in each model’s BIC and the mean BIC of all models examined. All hybrid models were comparable, but the model with the lowest BIC (a0b1s_hybrid, blue box) contained a single *α* positive learning rate, a single inverse temperature parameter, □, and a single strategy parameter,ξ_1_, that indicated the likelihood for an animal to stick with their previous choice if they had received a reward. The full winning model is written in the schematic in (A). (C) We generated data by simulating the model with parameters fit on individual sessions. Then, we inspected the group similarity between simulated data (blue dotted line for male *Tsc2* Het mice and green dotted line for female *Tsc2* Het mice) and found that the model simulated with fit model parameters captures averaged mouse learning curve data. (D-E) Examples of individual trial-by-trial model fits for both male (D) and female mice (E). The blue for males (D) and the green for females (E) indicate the fraction correct performance for session 1 for two example animals from each sex. The lighter, transparent line indicates the simulated data from the winning hybrid model and the thin black dashed line represents the inferred probability that the mouse is in the latent “engaged” state, i.e. following the RL policy, as opposed to the biased state. (F) There was no significant difference in normalized bias between male Het and WT groups. (G) There was a significant difference between *Tsc2* Het and WT males in learning rate *α*+ (plotted in log scale for visibility), Mann-Whitney test, *U* =54, *p* = 0.04, as well as *α*+ *□, a post-hoc variable that captures the overall impact of an outcome on the policy, (I) Mann-Whitney test, *U* = 41, *p* = 0.008. There was no difference in the □ parameter (H) in males. In females, there were no differences between *Tsc2* Het and WT mice in normalized bias (J), log(*α*)+ (K), □ (L), or log(*α*+* □) (M). Full statistical results are reported in the main text. Error bars represent SEM. \**p*<0.05, \*\**p*<0.01.

Our model comparison results, based on Bayesian Information Criterion (BIC), showed that hybrid models combining RL and biased policies significantly outperformed the RL component alone (Figure 2B, Wilcoxon signed-rank test: *z* = −3.31, *p* < 0.001 between a0b3s and a0b3s_hybrid; see Materials & Methods for detailed model specifications). Among the multiple hybrid models we evaluated, a0b1s_hybrid had the best fit to choice behavior (Wilcoxon signed-rank test: *z* = 2.10, *p* = 0.036 between a0b1s_hybrid and a0b3s_hybrid; *z* = 2.98, *p* = 0.0028 between a0b1s_hybrid and a0b2s_hybrid). The best fitting model (a0b1s_hybrid) included a single learning rate (*α*+), an inverse temperature parameter (□) that controls choice stochasticity, a single strategy parameter (ξ_1_) that measures “stickiness” (the likelihood of a mouse to repeat their previous action), latent state transition probabilities (lapse and recover), a bias parameter, and a fixed parameter set at 0 for learning from unrewarded trials. Model parameters were found to be identifiable (Supplemental Figure 2). At the group level, the hybrid model captured decision accuracy across learning (Figure 2C), which was remarkable given the highly variable trajectories observed in the first session (Figure 2D-E).

Using the best fit model, we compared parameters and variables across groups. In Session 1, male *Tsc2* Het mice exhibited a significantly higher alpha learning rate, *α*+, which signifies reliance on previous reward outcomes to guide future choices, compared to WT animals (Figure 2G: Mann-Whitney test, *U* = 54, *p* = 0.04). This significant effect was maintained in an examination of a post-hoc variable, log (*α* * □), that captures the incremental change in policy (Daw, 2011) (Figure 2I: Mann-Whitney test, *U* = 41, *p* = 0.008).

All other parameters were comparable between male *Tsc2* Het and WT mice. There was no difference between male *Tsc2* Het and WT mice in the normalized bias parameter, which measures the degree to which behavior is biased towards a specific choice (Figure 2F: Unpaired *t*-test, *t*(26) = 1.94, *p* = 0.06) or in the softmax □ parameter (Figure 2H: Mann-Whitney test, *U* = 66, *p* = 0.15) or ξ_1_ (Supplemental Figure 1L, Mann-Whitney test: *U* = 80, *p* = 0.43). The probability to transition from a biased to an engaged state (recover) and back (lapse) was also comparable between both *Tsc2* Het and WT mice (Supplemental Figure 1M-N: Mann-Whitney test, recover: *U =* 96, *p* = 0.96; lapse: *U* = 87, *p* = 0.65). Finally, using the fitted parameters including latent state transition probabilities, we calculated post-hoc each animal’s probability of being in an engaged state, “p(engaged)”, on each trial and found no difference in the mean p(engaged) value between male *Tsc2* Het and WT mice (Supplemental Figure 1O: Mann-Whitney test, *U* = 87, *p* = 0.65) or in the p(engaged) frequency distribution (Supplemental Figure 8A).

When female *Tsc2* Het and WT mouse data were fit with the same winning model, a0b1s_hybrid, we found no significant difference in log(*α*+) learning rate, the inverse temperature parameter □, or in the post-hoc variable, log(*α*+ * □) (Figure 2K-M: Mann-Whitney tests: *α*+: *U* = 43.5, *p* = 0.64; □, *U* = 34, *p* = 0.24; *α*+ * □: *U* = 42, *p* = 0.57). There was also no significant difference between female *Tsc2* Het and WT mice in the normalized bias parameter, indicating that neither Het nor WT mice exhibited a stronger bias towards a particular choice (Figure 2J: Mann-Whitney test, *U* = 46.5, *p* = 0.81). Additionally, no differences were observed in other fitted parameters or post-hoc calculations, including stickiness (ξ_1_), transition probabilities (recover and lapse), and mean p(engaged), which estimates the likelihood of staying in the engaged state (Supplemental Figure 1P-S: Mann-Whitney test, ξ_1_: *U =* 44, *p* = 0.66; recover: *U* = 44, *p* = 0.68; lapse: *U* = 40, *p* = 0.48; p(engaged): Mann-Whitney test, *U* = 40, *p* = 0.48).

### Early adolescent male *Shank3B* Het mice show a gain of function in early learning

*Shank3B* mice show corticostriatal hyperconnectivity in development (Peixoto et al., 2016; 2019), a phenotype that overlaps with observations in TSC models (Benthall et al., 2021). Therefore, we were interested in testing if there was a convergent reinforcement learning phenotype between *Tsc2* and *Shank3B* Hets. Adolescent male *Shank3B* Het mice (n = 12) along with WT littermates (n = 13) were water-restricted and trained in the odor-based 2AFC task as described (Materials and Methods). In the first odor learning session (Session 1), male *Shank3B* Het mice showed stronger performance (fraction correct) than WT littermates (Figure 3A: 2 way RM ANOVA: geno: *F*(1, 23) = 3.3, *p* = 0.08; time: *F*(1.96, 45.2) = 26.08, *p* < 0.0001; geno x time: *F*(3, 69) = 5.3, *p* = 0.002, with a significant post-hoc difference in quartile 4). ITI movement times were comparable between genotypes (Supplemental Figure 3A: Mann-Whitney test, *U* = 65, *p* = 0.50).

**Figure 3:**
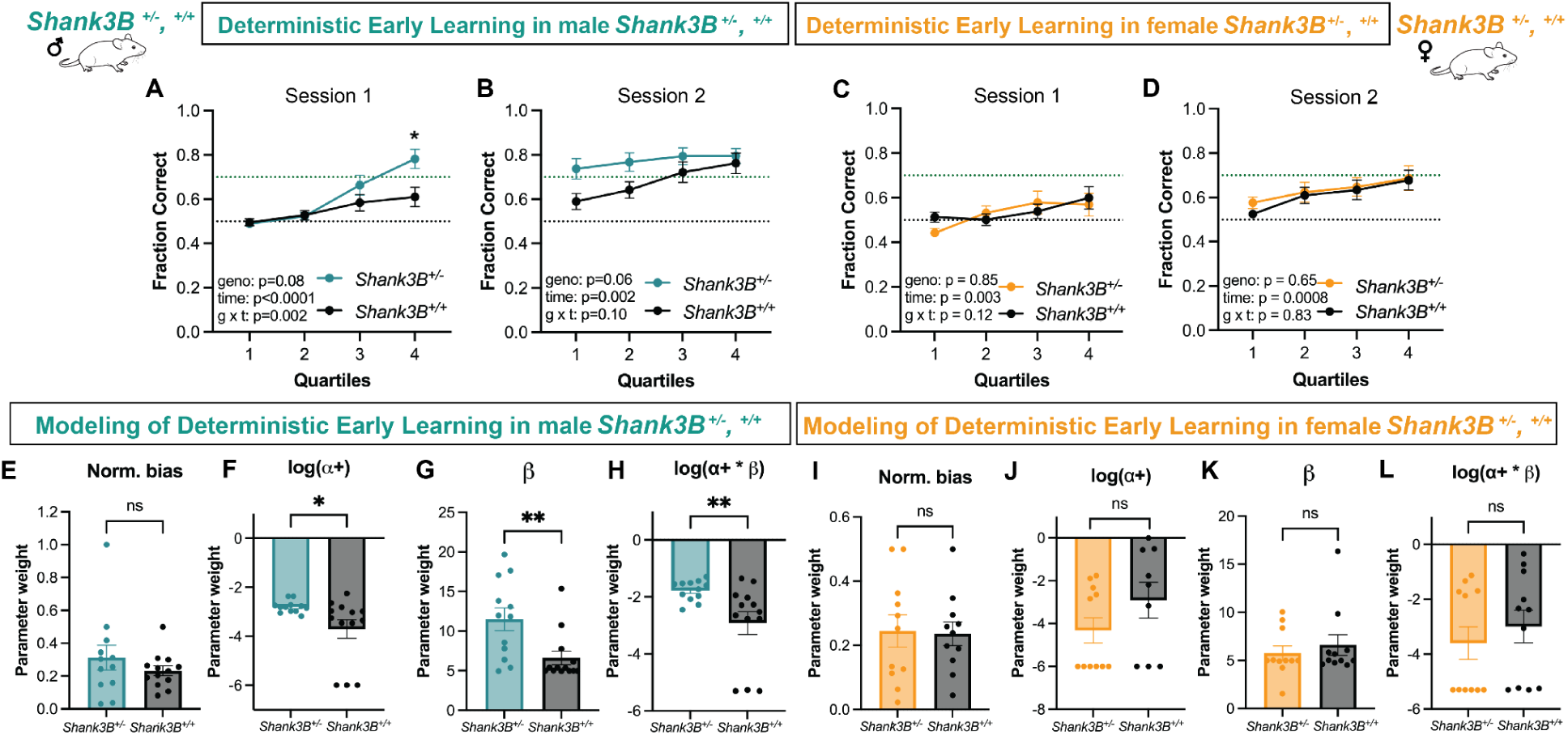
Developing male, but not female *Shank3B* Het mice show a gain of function in learning and a higher learning rate during early training in an odor based 2AFC task. (A) Male *Shank3B Het* showed stronger learning performance than WT littermates in the first training session, Session 1. This was captured by a significant main effect of time and genotype x time interaction. A Tukey post-hoc comparison test revealed a significant difference between Het and WT performance in the fourth quartile, indicated by the asterisk (p = 0.01). (B) There was a main effect of time but no genotype or genotype x time interaction in Session 2. We also tested female *Shank3B* Het and WT mice in the same odor-based 2AFC task on the same schedule. (C-D) Female Het and WT mice did not differ in learning performance for Session 1 or for Session 2 as evidenced by lack of significant main effect of genotype or an interaction. We next fit computational models to *Shank3B* male and female early learning data. Male *Shank3B* Hets and WT did not differ in normalized bias parameter (E), but did differ significantly in *α*+ (F) (plotted in log scale for visibility), □ (G), and the *α*+*□ variable (H). Female *Shank3B* Hets and WT did not differ significantly for any modeling parameter: (I) normalized bias parameter, (J) *α*+, (K) □, (L) *α*+*□. Full statistical results are reported in the main text. Male *Shank3B* Het, n = 12; Male *Shank3B* WT, n = 13; Female *Shank3B* Het, n = 11; Female *Shank3B* WT, n = 11. Error bars represent SEM. \**p*<0.05, \*\**p*<0.01.

In the second session, male *Shank3B* Het sustained their higher level of performance in the first quartile, but *Shank3B* WT improved and closed the gap (Figure 3B: 2-way RM ANOVA: geno: *F*(1, 23) = 3.73, *p* = 0.06; time: *F*(1.57, 36.19) = 7.46, *p* = 0.002; geno x time: *F*(3, 69) = 2.10, *p* = 0.10). ITIs in male *Shank3B* were again comparable (Supplemental Figure 3A: Mann-Whitney test, *U* = 56, *p* = 0.56). Performance showed a negative correlation with ITI that was statistically significant in *Shank3B* Het, but not WT (Spearman’s correlation: Het: *r* = −0.78, *p* < 0.0001, WT: *r* = −0.12, *p* = 0.48).

### Early adolescent female *Shank3B* Het mice have no gain of function in early learning

To test how haploinsufficiency of *Shank3B* affects females, we trained a separate cohort of adolescent female *Shank3B* Het (n = 11) and WT (n = 11) animals in the same odor-based 2AFC task. Female *Shank3B* Het and WT mice showed comparable performance in Session 1 (Figure 3C: 2-way RM ANOVA: geno: *F*(1, 20) = 0.03, *p* = 0.85; time: *F*(2.22, 44.44) = 6.26, *p* = 0.003; geno x time: *F*(3, 60) = 1.96, *p* = 0.12) and Session 2 (Figure 3D: 2-way RM ANOVA: geno: *F*(1, 20) = 0.20, *p* = 0.65; time: *F*(1.77, 35.43) = 9.45, *p* = 0.0008; geno x time: *F*(3, 60) = 0.29, *p* = 0.83).

Female *Shank3B* WT animal performance was comparable to male *Shank3B* WT performance in both sessions (Supplemental Figure 3B-C: Session 1: 2-way RM ANOVA: sex: *F*(1, 22) = 0.19, *p* = 0.66; time: *F*(2.00, 44.07) = 7.37, *p* = 0.001; sex x time: *F*(3, 66) = 0.61, *p* = 0.60; Session 2: 2-way RM ANOVA: sex: *F*(1, 22) = 2.13, *p* = 0.15; time: *F*(1.50, 33.05) = 12.82, *p* = 0.0003; *F*(3, 66) = 0.42, *p* = 0.73). Performance observed in male *Shank3B* Hets was higher compared to female *Shank3B* Hets with a significant main effect of sex and sex x time interaction in session 1 (Supplemental Figure 3D-E: Session 1: 2-way RM ANOVA: sex: *F*(1, 21) = 4.75, *p* = 0.04; time: *F*(2.12, 44.53) = 21.95, *p* < 0.0001; sex x time: *F*(3, 63) = 5.55, *p* = 0.001) and main effect of sex in session 2 ( Session 2: 2-way RM ANOVA: sex: *F*(1, 21) = 7.26, *p* = 0.01; time: *F*(1.83, 38.55) = 5.0, *p* = 0.01; sex x time: *F*(3, 63) = 0.48, *p* = 0.69).

Female Shank3B Het and WT animals showed no significant difference in ITIs in either the first or second odor learning sessions (Supplemental Figure 3F: Session 1: Mann-Whitney: *U* = 23, *p* = 0.07. Session 2: Unpaired *t*-test: *t*(15) = 1.87, *p* = 0.08). We observed a significant negative relationship between fraction correct (performance) and ITI in both Het and WT *Shank3B* females (Spearman’s correlation: *r* = −0.66, *p* < 0.0001; WT: Pearson’s correlation: *r* = −0.48, *p* = 0.002).

Male *Shank3B* Het mice had significantly faster ITIs on average compared to female Het mice (Session 1: *t*(19) = 4.54, *p* = 0.0002; Session 2: *U* = 15, *p* = 0.01). In contrast, mean ITIs were comparable between WT *Shank3B* male and WT *Shank3B* females (Session 1: *U* = 40, *p* = 0.13; Session 2: *U* = 26, *p* = 0.04). Last, we performed an additional analysis to test whether daily weight of male and female *Shank3B* Het and WT mice was correlated with performance. We found no significant correlation between these variables in Het or WT *Shank3B* mice (Supplemental Figure 3G-J).

### Gain of function is captured by higher learning rates in early adolescent male, not female, *Shank3B* Het mice

To explore latent structures in trial-by-trial learning, we next fit the *Shank3B* data using the set of models described above and detailed in our methods (Materials and Methods). Just as for the *Tsc2* mouse data, the winning model for *Shank3B* data was a0b1s_hybrid (Supplemental Figure 4A), which allowed for dynamic policy shifting between an RL-based “engaged” strategy and a biased, or “disengaged,” strategy (see Figure 2A). The model correctly captured decision accuracy both at the group level (Supplemental Figure 4B) and at the individual level (Supplemental Figure 4C) for the *Shank3B* data.

Like *Tsc2* Het males, *Shank3B* Het males also showed significantly higher *α*+ learning rates compared to WT littermates (Figure 3F: Mann-Whitney test *U* = 35, *p* = 0.01); this difference was also observed when examined with the post-hoc *α*+ * □ variable (Figure 3H: Mann-Whitney test *U* = 31, *p* = 0.009). Similar to *Tsc2* Hets, *Shank3B* Hets did not differ from WT animals in the normalized bias parameter, indicating that they did not prefer any particular side (Figure 3E, Mann-Whitney test, *U* = 61.5, *p* = 0.38), nor did they differ from WT in the strategy parameter, recover, lapse, or our post-hoc estimation of mean time spent in an engaged state (Supplemental Figure 4G-J: ξ_1_ : *U* = 56.5, *p* = 0.25; recover: *U* = 59; *p* = 0.31; lapse: *U* = 64, *p* = 0.46; p(engaged): Unpaired *t*-test, *t*(22) = 1.05, *p* = 0.30). *Shank3B* Hets and WT also did not differ in their distribution of p(engaged) frequency (Supplemental Figure 8B). However, different from the *Tsc2* male results, *Shank3B* Het males also had a significantly higher □ parameter than *Shank3B* WT males (Figure 3G: Mann-Whitney test, □: *U* = 30, *p* = 0.008), indicating that, in addition to learning rapidly from outcomes, they may also have a less noisy or exploratory policy than WT littermates.

Female *Shank3B* Het and WT mice were also fit with the same set of models. The winning model for *Shank3B* females was the same a0b1s_hybrid model (Supplemental Figure 4D) and was validated both in group learning curves (Supplemental Figure 4E) and individual learning trajectories (Supplemental Figure 4F). Female *Shank3B* Hets did not differ significantly from WT mice in any fit parameter or post-hoc calculation included in our comparisons (Figure 3I-L: normalized bias: Unpaired *t*-test, *t*(20) = 0.13, *p* = 0.89; *α*+: Mann-Whitney test, *U* = 54, *p* = 0.67; □: Mann-Whitney test, *U =* 55, *p* = 0.74; *α*+*□: Mann-Whitney test, *U =* 48, *p* = 0.43; Supplemental Figure 4K-N: ξ_1_: Mann-Whitney, *U* = 40.5, *p* = 0.19; recover: Unpaired *t*-test, *t*(20) = 0.35, *p* = 0.72; lapse: Mann-Whitney, *U* = 54, *p* = 0.69; p(engaged): *t*(20) = 0.53, *p* = 0.59).

### Gain of function in learning in *Tsc2* and *Shank3B* Het mice is reward schedule dependent

We next investigated whether altering the reward schedule could influence the enhanced performance and learning rates observed in *Tsc2* and *Shank3B* Het males. Past work has suggested that neurocognitive processes captured by RL models, such as *α* learning rates or □ values, are not simply intrinsic to the individual and instead, can vary depending on the context in which the learning takes place (Eckstein, Wilbrecht & Collins, 2021; Eckstein et al., 2022). This especially applies to learning across development (Wilbrecht & Davidow, 2024) and there is evidence that autistic individuals learn differently from probabilistic feedback (Koegel et al., 1979; Solomon et al., 2011; South et al., 2014; Lawson, Mathys, & Rees, 2017).

We trained new cohorts of *Tsc2* and *Shank3B* Het and WT mice (*Tsc2* Het, n = 10; *Tsc2* WT, n = 10; *Shank3B* Het, n = 11; *Shank3B* WT, n = 11) in the same odor-based 2AFC task, with one key difference: correct choices were unrewarded 10-20% of the time, introducing probabilistic outcomes (see Materials & Methods).

All mice were able to learn stimulus-action contingencies, despite the introduction of uncertainty into the environment. But, unlike the genotype differences observed in our first experiment, under a probabilistic reward schedule, task performance of male *Tsc2* Hets and male *Shank3B* Hets was comparable to their WT littermates in both Session 1 (Figure 4A: *Tsc2,* Session 1: 2-way RM ANOVA: geno: *F*(1, 18) = 0.47, *p* = 0.50; time: *F*(1.61, 29.1) = 15.3, *p* < 0.0001; geno x time: *F*(3, 54) = 0.13, *p* = 0.93; Figure 4D: *Shank3B*, Session 1: geno: *F*(1, 20) = 0.25, *p* = 0.61; time: *F*(1.87, 37.42) = 13.16, *p* < 0.0001; geno x time: *F*(3, 60) = 2.45, *p* = 0.07), and Session 2 (Figure 4B: *Tsc2,* Session 2: 2-way RM ANOVA: geno: *F*(1, 18) = 0.16, *p* = 0.68; time: *F*(2.01, 36.22) = 5.4, *p* = 0.008; geno x time: *F*(3, 54) = 0.25, *p* = 0.85; Figure 4E: *Shank3B*, Session 2: geno: *F*(1, 20) = 0.08, *p* = 0.77; time: *F*(2.15, 43.15) = 13.63, *p* <0.0001; geno x time: *F*(3, 60) = 2.29, *p* = 0.08).

**Figure 4:**
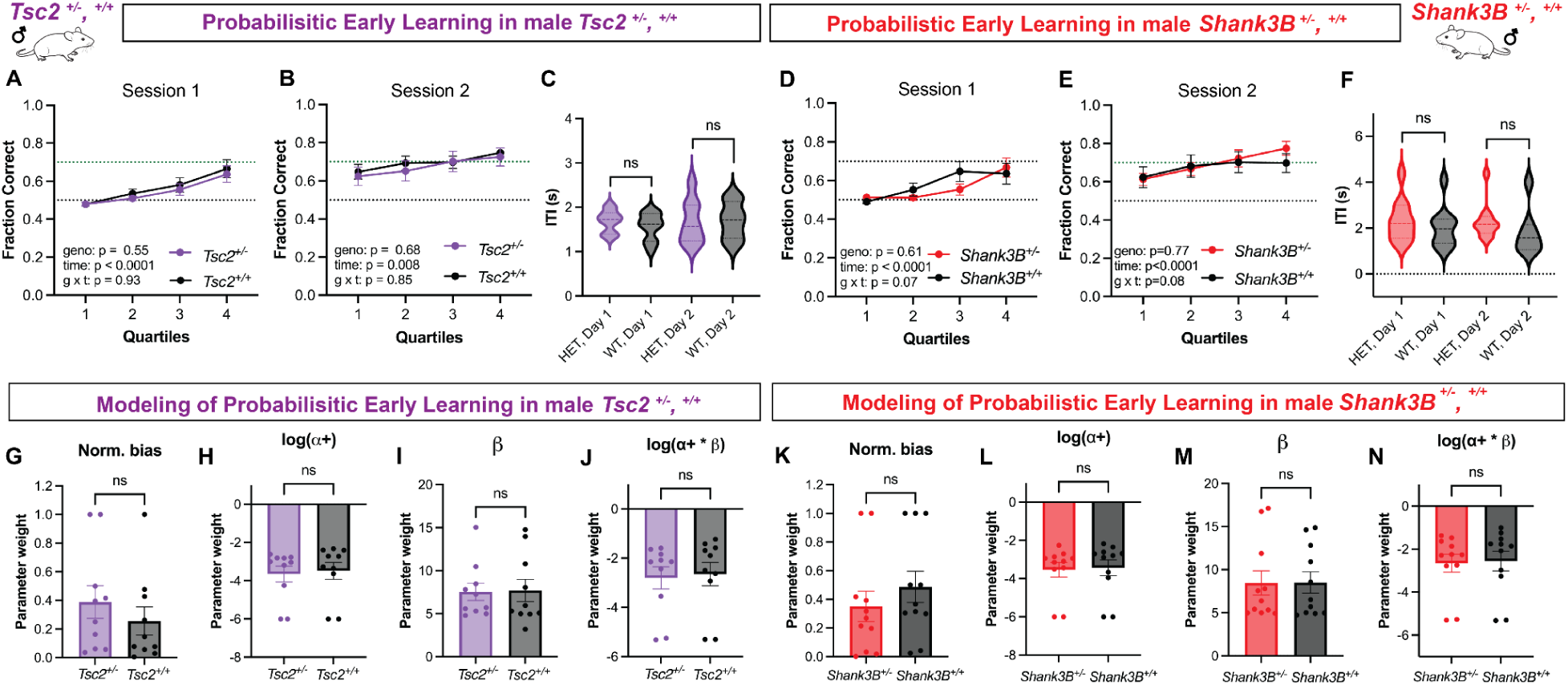
Gain of function in early adolescent male *Tsc2* and *Shank3B* Het is no longer apparent when reward schedule is probabilistic. We tested male *Tsc2* (A-C) and male *Shank3B* (D-F) in a probabilistic version of the odor-based 2AFC task where 10-20% of correct choices in session 1 and 2 were not rewarded (compared to Figures 1-3 data where 100% of correct choices were rewarded). Male *Tsc2* Het and WT showed comparable learning performance in this probabilistic context in both session 1 and 2 (A,B). ITI speeds were also comparable (C) between Day 1 (Het, n = 8; WT, n = 9) and Day 2 (Het, n = 10; WT, n = 9). (D,E) Similarly, in male *Shank3B* mice, Het and WT learning performance was comparable between genotypes for both sessions. (F) There was also no significant difference between groups in ITI for either Day 1 (Het, n = 11; WT, n = 10) or Day 2 (Het, n = 11; WT, n = 7). We next fit our series of models to probabilistic learning trial-by-trial data. We found that there were no significant differences between model parameters (G-J) in male *Tsc2* Het and WT mice nor between (K-N) male *Shank3B* Het and WT mice when performing in the probabilistic task context. Full statistical results are reported in the main text. *Tsc2* Het, n = 10; *Tsc2* WT, n = 10; *Shank3B* Het, n = 11; *Shank3B* WT, n = 11. Error bars show SEM. Violin plots show quartiles (25% and 75%), mean, and range.

ITIs in Sessions 1 and 2 were also comparable between Het and WT mice from both lines in the probabilistic schedule. In Tsc2 animals (Figure 4C), ITIs did not differ between groups in Session 1 (Unpaired *t*-test: *t*(18) = 1.03, *p* = 0.31) or Session 2 (Unpaired *t*-test: *t*(14) = 0.08, *p* = 0.93). In Shank3B animals (Figure 4F), ITIs were also comparable in Session 1 (Mann-Whitney test: *U* = 42, *p* = 0.38) and showed a non-significant trend in Session 2 (Mann-Whitney test: *U* = 19, *p* = 0.08).

### Male *Tsc2* and *Shank3B* Het mice have comparable learning rates to WT males when trained with a probabilistic reward schedule

We next fit our family of RL models to the data from early adolescent males trained in the probabilistic context. Despite a lack of difference between genotypes in basic performance, it is still possible that there could be differences in latent parameters. We found that the winning model (a0b1s_hybrid) for *Tsc2* and *Shank3B* males trained in the probabilistic context was the same winning model as for the deterministic contexts (Supplemental Figure 5A,D) and was validated at both the group level (Supplemental Figure 5B,E) and for individual trajectories (Supplemental Figure 5C,F).

When trained in the probabilistic context, *Tsc2* Het and WT male mice had statistically comparable parameter values (Figure 4G-J: normalized bias: Mann-Whitney, *U* = 38, p = 0.39; *α*+: Mann-Whitney test, *U* = 41, p = 0.51; □: Mann-Whitney test, *U* = 45, *p* = 0.73; *α*+*□: Mann-Whitney test, *U* = 44, *p* = 0.68; Supplemental Figure 5G-J: ξ_1_: Unpaired *t*-test, *t*(18) = 1.9, *p* = 0.07; recover: Mann-Whitney test, *U* = 32, *p* = 0.18; lapse: Mann-Whitney test, *U* = 31, *p* = 0.16; mean p(engaged): Mann-Whitney test, *U* = 31, *p* = 0.16).

Similarly, *Shank3B* Het and WT male mice trained in the probabilistic context also showed statistically comparable parameter values (Figure 4K-N: normalized bias: Mann-Whitney, *U* = 43, *p* = 0.25; *α*+: Mann-Whitney test, *U* = 48, *p* = 0.42; □: Mann-Whitney test, *U* = 58, *p* = 0.89; *α*+*□: Mann-Whitney test, *U* = 55, *p* = 0.74; Supplemental Figure 5K-N: ξ_1_: Unpaired *t*-test, *t*(20) = 1.03, *p* = 0.31; recover: Unpaired *t*-test with welch’s correction, *t*(14.08) = 1.67, *p* = 0.11; lapse: Mann-Whitney test, *U* = 45, *p* = 0.33; p(engaged): Mann-Whitney test, *U* = 54, *p* = 0.69). These data suggest that the gain of function in learning rate observed in male *Tsc2* and *Shank3B* Het mice compared to WT was sensitive to uncertainty in reward schedule.

### A subset of male *Shank3B* KO exhibit high trial numbers and gain of function in learning

While *Shank3B* Het mice more closely reflect the mutations reported in humans (Leblond et al., 2014; Cammarata-Scalisi et al., 2022), many animal studies focus on *Shank3B* KO mice (Mei et al., 2016; Jaramillo et al., 2016; Wang et al., 2017; Dhamne et al., 2017; Rendall et al., 2019). Thus, we also chose to train developing *Shank3B* KO male mice in the deterministic odor-based 2AFC task (100% reward).

We found that, on average, developing male *Shank3B* KO (n = 13) had comparable performance to *Shank3B* WT (littermates whose data was shown above) in the first session (Figure 5A: 2-way RM ANOVA: genotype: *F*(1, 24) = 1.6, *p* = 0.21; time: *F*(2.01, 48.24) = 16.92, *p* < 0.0001; genotype x time: *F*(3, 72) = 1.11, *p* = 0.34) and the second session (Figure 5B: 2-way RM ANOVA: genotype: *F*(1, 25) = 0.91, *p* = 0.34; time: *F*(1.63, 40.94) = 7.2, *p* = 0.003; geno x time: *F*(3, 75) = 1.13, *p* = 0.33).

**Figure 5:**
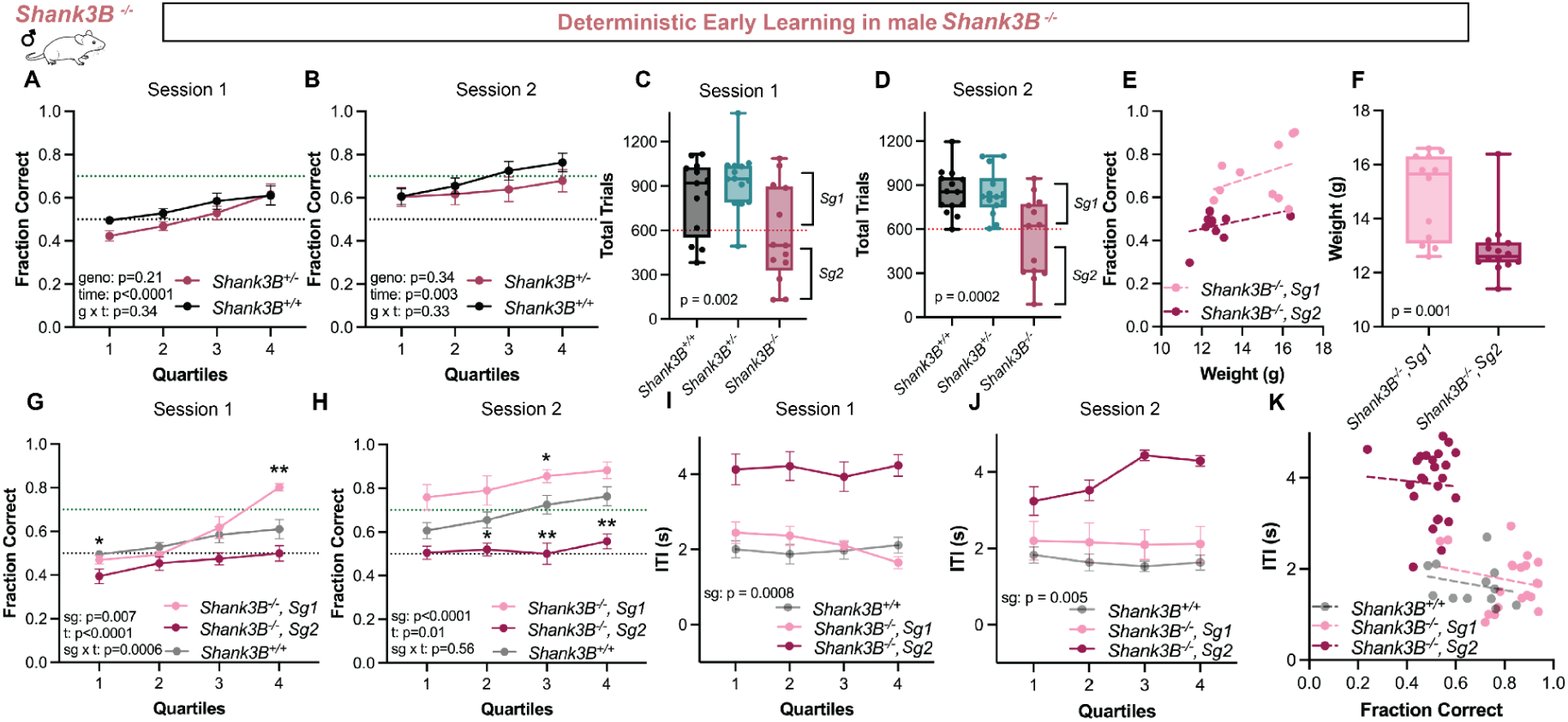
Early adolescent male *Shank3B* KO mice exhibit bimodal levels of trial completion and the more active subgroup shows gain of function in learning performance. (A-B) In Session 1 and Session 2 the full group of male *Shank3B* KO mice (n = 13) had comparable learning performance to male *Shank3B* WT animals (n = 14). (C,D) Looking at total trials completed in each session, we noted that the total number of completed trials was bimodal for KO mice. Total trials completed were comparable to WT and Het siblings for one subgroup of KO (n = 5) (Sg1), but there was also a second subgroup (Sg2) (n = 8) that completed substantially fewer trials per session. Box and whisker plots show 25%, mean, 75%, and range with individual points indicating total number of trials for each individual mouse. Examining these groups more closely, revealed that (F) Sg1 weighed significantly more than Sg2, but (E) weight was not correlated with fraction correct performance in either subgroup. Based on these differences we decided to split the KO group into two subgroups and compare their learning to WT. (G-H) Sg1 KOs, who completed a typical range of trials, showed significantly better learning performance than WT in the odor based 2AFC in both Session 1 and Session 2. Sg2 KOs, who completed substantially fewer trials, performed at chance levels, significantly lower than WT in both sessions. Asterisks indicate post-hoc comparisons of subgroups to WT. In Session 1, quartile one, Sg2 KOs had significantly lower performance than WT (*p* = 0.03) and in quartile four, Sg1 KOs had significantly higher performance than WT (*p* = 0.002). In Session 2, Sg2 KOs had significantly lower performance than WT in quartile two (*p* = 0.01), three (*p* = 0.007), and four (*p* = 0.002). Sg1 outperformed WT in quartile three (*p* = 0.04). (I-J) Examination of intertrial interval (ITI) movement times showed that Sg2 was also significantly slower to move between ports than Sg1 in both sessions and that Sg1 ITI movement times more closely resembled WT mice, including in their correlation with fraction correct performance (K). Violin plots show quartiles (25% and 75%), mean, and range. Error bars represent SEM. \**p*<0.05, \*\**p*<0.01, \*\*\**p*<0.001, \*\*\*\**p*<0.0001.

In our experience, most developing mice, WT or otherwise, complete well over 600 trials in each behavioral session. However, in examining total trials completed by individual *Shank3B* KO mice across each session, we observed a bimodal distribution in the number of trials completed by *Shank3B* KO mice (see Supplemental Figure 6A-B). The first subgroup of KO animals (n = 5) (Sg1) showed typical numbers of trial completed in a session (compared to other *Shank3B* genotypes), with 840.5 ± 125.50 (SD) total trials per session (*Shank3B* Het averaged 890.21 ± 100.36 (SD) and *Shank3B* WT averaged 841.29 ± 110.45 (SD)). In contrast, the second subgroup of KO mice (n = 8) (Sg2) completed only 310.46 ± 160.55 (SD) trials, on average, per AB session (Figure 5C-D). *Shank3B* KO subgroups came from multiple breeder pairs and individuals from the same litter could be found in the two subgroups.

When we compared the task performance of these two *Shank3B* KO subgroups to WT, we found that subgroup 1 significantly out performed WT whereas subgroup 2 showed a deficit compared to WT in both sessions (Figure 5G: Session 1, 2-way RM ANOVA: subgroup: *F*(2, 23) = 6.12, *p* = 0.007; time: *F*(2.07, 47.68) = 26.93, *p* < 0.0001; subgroup x time: *F*(6, 69) = 4.57, *p* = 0.0006; asterisks indicate Dunnett post-hoc tests comparing subgroup vs WT for that quartile) (Figure 5H: Session 2: subgroup: *F*(2, 23) = 14.14, *p* < 0.0001; time: *F*(1.62, 37.32) = 5.00, *p* =0.01; subgroup x time: *F*(6, 69) = 0.80, *p* = 0.56; asterisks indicate Dunnett post hoc tests comparing subgroup vs WT for that quartile). Further analysis revealed Sg1 KO mice weighed more than the Sg2 KO mice (Figure 5F: Mann-Whitney test: *U* = 4, *p* = 0.001), but weight was not correlated with higher performance within either group (Figure 5E: Pearson’s correlation: Sg1: *r* = 0.36, *p* = 0.29; Sg2: *r* = 0.36, *p* = 0.26). Sg1 KO mice also had faster ITIs in both sessions (Figure 5I-J, Session 1: Unpaired *t-*test, *t*(11) = 4.59, *p* = 0.0008; Session 2: Unpaired *t*-test, *t*(10) = 7.37, *p* < 0.0001). ITIs from Sg1 KOs were comparable to both *Shank3B* Het and *Shank3B* WT littermates (One-way ANOVA: Session 1: *F*(2, 27) = 0.95, *p* = 0.39; Session 2: *F*(2, 25) = 0.14, *p* = 0.86).

In sum, our analysis of male *Shank3B* KO subgroups suggests that a subset of these mice show deficits in all of our metrics, while another subset that appears more typical in basic metrics shows a gain of function in reinforcement learning, similar to *Shank3B* Het and *Tsc2* Het mice.

## Discussion

In the present study, we used an odor-based 2AFC task and reinforcement learning models to examine learning in mice with mutations in two autism risk genes, *TSC2* and *SHANK3*. We chose these genes for two reasons. First, single gene mutations in *TSC2* or *SHANK3* in humans cause syndromes that are associated with autism diagnoses and together, may account for approximately 1% of ASD cases (de la Torre-Ubieta et al., 2016). Second, mice with mutations in *Tsc1*, *Tsc2*, or *Shank3B* at young ages have been shown to have an increase in corticostriatal connectivity (Peixoto et al., 2016; Benthall et al., 2018). This hyperconnectivity is potentially linked to enhanced motor learning in TSC lines (Benthall et al., 2021; Cording and Bateup, 2023) and grooming behaviors in mice that have parallels to repetitive, restricted behaviors in humans (Wang et al., 2017). Using an odor-based 2AFC, we found a convergent gain of function in the first days of odor learning for both adolescent male *Tsc2* Het and *Shank3B* Het mice that was not observed in females and was dependent on a deterministic (100%) reward schedule. When we analyzed trial-by-trial learning using an RL-based mixture model that accounts for periods of task disengagement, we found that the gain of function in early learning performance in both *Tsc2* and *Shank3B* males was driven by a significant difference in the *α*+ learning rate parameter – a difference that was not observed in females. This gain of function in learning rate in male Hets was no longer apparent when *Tsc2* and *Shank3B* males were trained with a probabilistic schedule, suggesting the learning phenotype was modulated by environmental uncertainty.

It may seem surprising that we see a gain of function in mice with mutations in autism risk genes. While it is true that previous studies of rodents with mutations in autism risk genes, including both *Tsc2* and *Shank3*, have shown deficits in hippocampal-mediated, perceptual, and social learning (Ehninger et al., 2008; Copping et al., 2017; Dhamne et al., 2017), there is a large literature showing a gain of function in motor learning in mice with mutations to a variety of autism risk genes, including *Tsc2* Hets (Penagarikano et al., 2011; Rothwell et al., 2014; Hulbert et al., 2020; Benthall et al., 2021; rev’d in Cording & Bateup, 2023; Cording et al., 2024). Gains in motor learning have been found using the accelerating rotarod, a task that engages the dorsal and ventral striatum (Cording & Bateup, 2023). One study found that a conditional knockout of *Tsc1* in direct pathway neurons of the dorsolateral striatum reduced endocannabinoid dependent LTD onto these neurons, enhanced their cortical inputs, and was sufficient to drive a gain of function in the rotarod (Benthall et al., 2021).

In *Shank3B* mice, the rotarod data are more complex as they vary by genotype (Het vs KO) and may also depend on the age of testing. Briefly, in adult *Shank3B* Het mice, rotarod performance has been found to be comparable to WT in two studies (Rendall et al., 2019; Cerilli et al., 2024). In *Shank3B* KO mice one study reports a day 1 gain of function then WT and Het “catch up” (Rendall et al., 2019) while another reports an overall loss of function compared to HET and WT (Cerilli et al., 2024). Notably, these rotarod tests were performed in adulthood when corticostriatal circuits may weaken in *Shank3B* mice (Peixoto et al., 2019). Future work should examine rotarod performance across different ages in *Shank3B* mice to examine if it is developmentally regulated.

In humans, ASD related restricted and repetitive behaviors have been correlated with changes in connectivity and structure of the dorsal striatum (Cerliani et al., 2015; Schuetze et al., 2016). It has been proposed that an increase in innate repetitive grooming behaviors in adult *Shank3B* KO mice may model these symptoms. Interestingly, this enhanced grooming in *Shank3B* KO has also been linked to disrupted endocannabinoid-dependent regulation of cortical inputs onto spiny projection neurons in the dorsal striatum (Wang et al., 2017) which has also been reported in studies of *Tsc1* (Benthall et al., 2021). In a behavioral task more similar to current study, a gain of function in perceptual discrimination learning has also been reported in *Shank3B* Hets and KO that is limited to the initial days of training (Rendall et al., 2019).

While rotarod learning, repetitive grooming, perceptual discrimination, and 2AFC learning look very different in practice, they all rely on the striatum and corticostriatal circuits (Balleine, Delgado, & Hikosaka, 2007; Lee et al., 2015; Znamenskiy et al., 2013; Cox & Witten, 2019; Tang et al., 2022). Building from this literature, we support and extend a working model in which enhanced corticostriatal connectivity (generated by diverse mechanisms), can facilitate a gain of function that can manifest in rotarod learning (Benthall et al., 2021; Cording & Bateup, 2023), repetitive behaviors like excess grooming (Wang et al., 2017), and 2AFC learning with an enhanced *α* learning rate (data presented here). The fact that two very different ASD risk genes produced a similar gain of function in early 2AFC learning in males, supports the models which suggest ASD may emerge from a convergent change in circuit function, possibly at the level of corticostriatal synapses, downstream of a variety of genetic changes (Fucillo, 2016; Cording and Bateup, 2023; Cording et al., 2024).

The fact that significant behavioral phenotypes and RL parameter differences were observed in males, but not females in both lines, also suggest convergent sex differences. This aligns with the human literature, where a male bias in autism diagnosis is commonly found (Maenner et al., 2023). Among those diagnosed with autism, sex differences in phenotypic expression are well documented, with males more often exhibiting pronounced restricted and repetitive behaviors than females (Hartley & Sikora, 2009; Mandy et al., 2012; Schuck, Flores, & Fung, 2019). In rodent models of ASD, similar sex differences have been reported. For instance, in some pharmacological models male rodents display more severe behavioral differences than females, in repetitive behaviors and tests of social behavior (Murta et al., 2023). Likewise, for multiple genetic models of autism, male mice show more putative autism-related phenotypes compared to females (Ferri et al., 2021). However, there are also studies of mice with mutations to other Autism risk genes that find significant phenotypes present in both males and females (Murta et al., 2023).

Our findings strongly support a role for sex in the expression of learning related phenotypes, but they do not help us isolate when genes and/or hormonal influences were critical to the generation of these differences. We collected data at P30-P35, which is just before the time of first estrus in female C57 Bl/6 mice reared in our lab (Piekarski et al., 2017). We did not monitor pubertal milestones or gonadal hormone levels in our mice, but hormones may impact learning and decision making (Delevich et al., 2021). Phenotypes may also fluctuate with age. Further research will be needed to better understand how developmental stage may interact with ASD genetics to influence phenotypes (Dahl et al., 2018; Nussenbaum & Hartley, 2019; Corbett et al., 2020; Muscatello et al., 2022).

There are of course numerous differences between human and mouse brains, but reinforcement learning in a 2AFC task may enable us to isolate latent variables comparable across species (Chase et al., 2024; Master et al., 2020). Recent research indicates that learning rates are not necessarily intrinsic to an individual but instead are shaped by interactions between the individual and the context of a learning environment (Nussenbaum & Hartley, 2019; Eckstein, Wilbrecht & Collins, 2021; Eckstein et al., 2022; Heald, Wolpert, & Lengyel, 2023). Our probabilistic schedule experiments were intended to probe the gain of function in *Tsc2* and *Shank3B* Het males by using a new cohort and training them with a different reinforcement schedule in the 2AFC task where correct choices were not rewarded 10-20% of the time. With this contextual change that increased uncertainty, we no longer observed a gain of function in performance or *α* learning rate in male *Tsc2* and *Shank3B* Hets compared to their WT littermates. From a bird’s eye perspective, reducing the reinforcement schedule could be seen as a phenotypic “rescue” that could be useful for training and interventions where enhanced learning may be disadvantageous. These data may be consistent with data from human participants that found changes in reinforcement schedule led to appearance and/or disappearance of performance differences between autistic and neurotypical participants (Solomon et al., 2011; Lawson, Mathys, & Rees, 2017; Koegel, 1972).

In conclusion, our data suggest diverse genetic changes associated with autism can interact with sex to lead to convergent behavioral outcomes. While these behavioral phenotypes stand out in males, they are still sensitive to environmental context and experience. Future studies may find there are alternate manifestations in females and that age and/or stage of testing also plays a large role. Human learning profiles may also differ from mice. One takeaway is that learning proceeds differently with ASD-related biology and may be sculpted by different rules. These observations support the celebration of some strengths in autism and may inform the design of future interventions and outcome metrics.

## Materials and Methods

### Animals

Males and females from two genetic lines were used for this experiment: *Tsc2* (JAX, #004686) and *Shank3B* (Jackson Laboratories Repository (JAX), #017688). We bred all experimental animals in-house, using HET and WT pairs for *Tsc2* and HET pairs for *Shank3B*. Breeders were maintained in clean cages by experienced researchers and cage changes were avoided while pups were small (under postnatal (P) day 10) to minimize disruption to care. All mice used in experiments were group housed in the animal facility under a reverse light/dark cycle (12h:12h) and were tested in the dark (active phase). Mice were habituated to chambers starting at P26-29 and were tested in development between P30-35. The following numbers of mice were used: male *Tsc2^+/-^*, n = 25; male *Tsc2^+/+^*, n = 23; female *Tsc2^+/-^*, n = 10; female *Tsc2^+/+^*, n = 10; male *Shank3B*^-/-^, n = 13; male *Shank3B^+/-^*, n = 23; male *Shank3B^+/+^*, n = 24; female *Shank3B^+/-^*, n = 11; female *Shank3B^+/+^*, n = 11. All procedures were approved by University of California, Berkeley Institutional Animal Care and Use Committee (IACUC).

### Odor-based Reinforcement Learning Task

Custom-built operant chambers that have an initiation port in the center and two lateral ports (Figure 1A) were used for 2AFC behavioral testing. Each port contains an infrared photodiode/phototransistor pair to accurately time port entry and exit (Island Motion, Tappan, NY). Side ports release water through valves (Neptune Research) and were regularly calibrated to ensure that exactly 2 µL of water was delivered for correct choices. Constant air flows through the center port at ∼0.5L/min and is redirected through one of the two possible odorant filters (Target2 GMF F2500-19) at trial initiation. The task was controlled using custom-built MATLAB scripts that also store outcome and action information.

Inexperienced early adolescent (postnatal day 27) male and female mice were first water-restricted and pre-trained for four days to initiate trials via nose poke to the center port and next to associate side ports with water reward. Next, animals entered “Session 1” where they first encountered an in-task odor pair: A (cinnamon: McCormick & Co, Hunt Valley, MD) and B (vanilla: McCormick Culinary, Hunt Valley, MD), which stably predicted the left and right port respectively for reward. Each session lasted 6-12 hours. Animals were all weighed at the end of each session and, if they did not gain weight within the session and/or across sessions, they were given supplemental water. All animals gained weight throughout their training (Supplemental Figure 1A) at a similar developmental trajectory to WT littermates (Supplemental Figure 7) and fraction correct performance was not correlated with weight in males or females (Supplemental Figure 1H-K; Supplemental Figure 3G-J). The training protocol was identical for the probabilistic reward schedule cohorts, except that 10-20% of correct choice trials were unrewarded. All mice experienced 10% unrewarded trials for Session 1 (90-0 protocol) and most mice also experienced 10% unrewarded trials for Session 2. A small subset of mice (3 *Tsc2* WT, 2 *Tsc2* Het, and 1 *Shank3B* WT) experienced both an 90-0 and 80-0 protocol on Session 2. No animals in the probabilistic cohort experienced deterministic rewards in the odor task prior to testing.

### Data and Statistical Analysis

Behavioral data was collected in MATLAB. The number of trials completed in each session was determined by individual mice, so each session for each mouse was divided into quartiles to examine the mean fraction correct. Intertrial interval (ITI) was calculated as time between side port exit and center port entry for each quartile. Fraction correct and ITI data for each group are shown as arithmetic mean and standard error of the mean (SEM) for each quartile in each session. Software used included Jupyter notebook, Python 3 and GraphPad Prism 10.

We checked assumptions of normality and variance using Shapiro-Wilk and Anderson-Darling tests (Graphpad Prism 10) and then used repeated measures 2-way analysis of variance (RM 2-way ANOVA) to examine performance over time in a session, Unpaired *t*-tests or Mann-Whitney tests for single time point comparisons, and Pearson and Spearman for tests of correlation (Graphpad Prism 10 or Jupyter Notebook Python 3). The significance level was set to *p* < 0.05 for all statistical tests.

Exclusion of some data was necessary. We strived to include all task behavior in calculations of fraction correct and trial by trial modeling, but had to exclude a small amount of trials where the mouse nose pokes were not sufficient to properly initiate or complete a trial (missed or aborted trials). To better reflect task-related behavior but to remove grooming sessions or other off-task behavior, we also excluded ITIs that were longer than 10 seconds (typically less than 5% of trials for each animal). Due to errors in data conversion, ITI data was missing for 22 of 308 total sessions. These sessions were excluded from ITI analysis and n-values reported in the figure legends reflect the number of mice for which all ITI data were available.

### Computational Modeling

#### Model specifications

We evaluated five RL-based models with variants of learning and decision-making mechanisms. All models learn by tracking and updating the values of both actions trial-by-trial. On any trial *t*∈[1, *T*], we denote the state (odor identity) as *s*_*t*_, action (go left or right) as *a*_*t*_ ∈ {*L*, *R*}, magnitude of reward as *r*_*t*_ ∈ {0, 1}, and value of an action *a* in state *s* as *Q*(*s*, *a*). On trial *t*, the model samples an action according to π_*t*_, a softmax policy that returns the probability of sampling the “go left” action given the current state *s*_*t*_ :

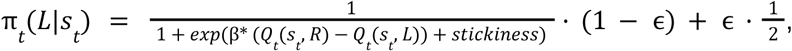

where β is the softmax inverse temperature parameter and ɛ parameterizes uniform decision noise. Stickiness = ξ · 𝟙_*a*_*t*−1_=*L*_ is a choice kernel defined by a set of parameters ξ (varying across models) that models the extent to which the animal sticks to the previously chosen action conditioned on the outcome of the previous action (i.e., *r*_*t*−1_) and whether the current odor is the same as the previous odor (i.e., *s*_*t*_ = *s*_*t*−1_); see below for further details on the “stickiness” kernels. When the chosen action *a*_*t*_ is rewarded, the action value is updated incrementally with a reward prediction error, following the standard delta rule:

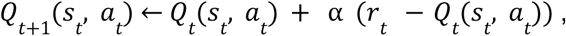

where α is the learning rate parameter, whose value depends on whether the chosen action *a*_*t*_ is rewarded. If it is rewarded, the learning rate takes on the value α_+_, whereas if the chosen action is unrewarded, the learning rate is α_−_.

Our baseline models include a standard RL model and the winning model in Chase et al. (2024), named RL_epsilon and a0b3s, respectively. RL_epsilon has five parameters: positive and negative learning rates (α_+_ and α_−_), softmax inverse temperature (β), uniform decision noise (ɛ), and one-back choice stickiness independent of states and outcomes (ξ). The a0b3s model also includes five parameters: α_+_, β, and three choice stickiness parameters conditioned on the outcome of the previous action and whether the state has changed from the previous trial to the current trial (ξ_1_ if the previous outcome is positive, ξ_2_ if the previous outcome is negative and the current state differs from the previous state, and ξ_3_ if the previous outcome is negative and the current state is the same as the previous state). Following Chase et al. (2024), when the chosen action is not rewarded, no action values are updated (i.e., the learning rate for negative rewards α_−_ is zero).

Extending the a0b3s model using dynamic noise estimation (Li et al., 2024), we tested three additional models with hybrid policies between a latent RL policy state and a latent biased policy state: a0b3s_hybrid, a0b2s_hybrid, and a0b1s_hybrid. We named our models based on the parameters in the RL policy: the “a0-” prefix indicates that the positive learning rate α_+_ is a free parameter while the negative learning rate α_−_ is fixed to 0, “-b-” denotes the softmax inverse temperature parameter β, and the “-3s”, “-2s”, and “-1s” suffixes specify the number of stickiness parameters. Specifically, a0b3s_hybrid includes the a0b3s model as a policy state, a0b2s_hybrid only includes ξ_1_ and ξ_2_ (fixed ξ_3_ = 0), and a0b1s_hybrid only includes ξ_1_ (fixed ξ_2_ =0 or ξ_3_ = 0).

The biased policy is a heuristic (i.e., not learned in the task) defined by a “bias” parameter between 0 and 1, which quantifies the degree of bias for the left action:

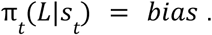

The hybrid model uses a hidden Markov model to construct a mixture policy between the RL and biased policies. It assumes the existence of two discrete hidden latent states (engaged or biased), which determine whether the agent follows the engaged RL-based policy or the biased heuristic policy. The transition probability from an engaged to a biased (lapse) and from a biased to an engaged state (recover) is described in Figure 2A, is equal to the transition probability from the current state into the same state, and can be summarized with the below relationship:

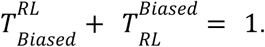

Its likelihood is estimated by extending the dynamic noise estimation framework described in (Li et al., 2024).

### Model fitting

All models were fitted to Session 1 with maximum a posteriori (MAP) estimation using a normal prior on the softmax inverse temperature parameter β with a mean of 5 and a standard deviation of 7, and uninformative priors for all other parameters. We used a prior for the β parameters as it is known to otherwise trade off with the learning rate parameter at fitting (Daw, 2011). All models were fit at the individual level.

### Model comparison

The fitted models were compared using the Bayesian Information Criterion (BIC) (Schwarz, 1978). BIC Figures can be found in Figure 2 for *Tsc2* mice as well as Supplementary Figure 4 and 5 for *Shank3B* mice and probabilistic training schedules, respectively. The best-fit parameters and latent variables were compared at the group-level for interpretation. We report the log-transformed values of the learning rates due to small values. The bias parameter is normalized by |*bias* − 0. 5| to indicate the un-sided magnitude of bias in choices. We additionally report *log*(α_+_ * β), a composite index between the positive learning rate and softmax inverse temperature, as a measure of the incremental change in policy that accounts for both the rate of learning updates and the stochasticity in decision-making (Daw, 2011).

### Model validation

To validate the model, we simulated choice behavior for each animal repeatedly for 1000 times using the MAP parameters obtained from model-fitting. For simulations with hybrid models, we used the latent state probability—P(RL) and P(Biased)—trajectories inferred from real data to estimate latent state occupancy. To validate how well the models captured behavior, we compared learning curves between these model simulations and the animal data that the models were fitted to. Group and individual validation curves can be found in Figure 2 for *Tsc2* mice as well as Supplementary Figure 4 and 5 for *Shank3B* mice and probabilistic training schedules, respectively.

### Parameter recovery

We performed parameter recovery analysis as a robustness check of the winning model, a0b1s_hybrid. To that end, we simulated choice data for each animal using the best-fit parameter values as ground-truth given the model-estimated latent state occupancy trajectory based on animal data. We then fitted the model back to the simulated data to obtain best-fit parameters, which were compared to the ground-truth parameter values (Supplementary Figure 2).

### Quantification of group-level p(engaged) distribution differences

To assess differences in the distributions of model-estimated engagement probability (p(engaged)) between groups (e.g., WT and Hets), we employed Kullback-Leibler (KL) divergence as a measure of difference between two probability distributions. KL divergence quantifies the extent to which one distribution diverges from another, with values close to zero indicating high similarity. Group-level p(engaged) distributions were constructed by dividing trial-level probabilities into 10 bins for each animal and averaging across animals within each group.

For statistical comparison, we performed a permutation test. Animal-level *p(engaged)* distributions were randomly reassigned to animals across groups, and KL divergence was recalculated for each permutation (10,000 iterations). The observed KL divergence between the original group distributions was compared to the distribution of permuted KL divergences to compute a p-value, defined as the proportion of permuted KL values greater than or equal to the observed value. This procedure provided a non-parametric assessment of whether observed group-level differences were significant. Error bars represent the standard error of the mean (SEM) for binned *p(engaged)* values.

## Acknowledgements

We thank Tory Benson, Anna Jahng, Katrina Manaloto, and Katrina Wong for their assistance in animal behavior and handling. We thank Kristen Delevich for helping to initiate these experiments and Helen Bateup, Dan Feldman, Rui Peixoto, Jonathan Tarbox, Katie Cording, and Katie Benthall for discussion.

This work was supported by a Pilot grant from the Simons Foundation Autism Initiative (SFARI) Award #613972 to L.W. and the National Institutes of Health (R01MH134514) to L.W.

## Author Contributions

**Juliana Chase**: Investigation, Conceptualization, Data Curation, Methodology, Writing - Original Draft, Visualization, Supervision, Analysis, Validation; **Jing-Jing Li**: Methodology, Software, Analysis, Investigation, Data Curation, Visualization, Writing; **Wan Chen Lin**: Software, Analysis; **Lung-Hao Tai**: Methodology, Software; **Fernanda Castro**: Investigation; **Anne GE Collins**: Resources, Supervision, Conceptualization, Funding acquisition, Writing - Editing; **Linda Wilbrecht:** Project Administration, Funding acquisition, Conceptualization, Supervision, Writing - Editing

## Supplemental Information

**Supplemental Figure 1:**
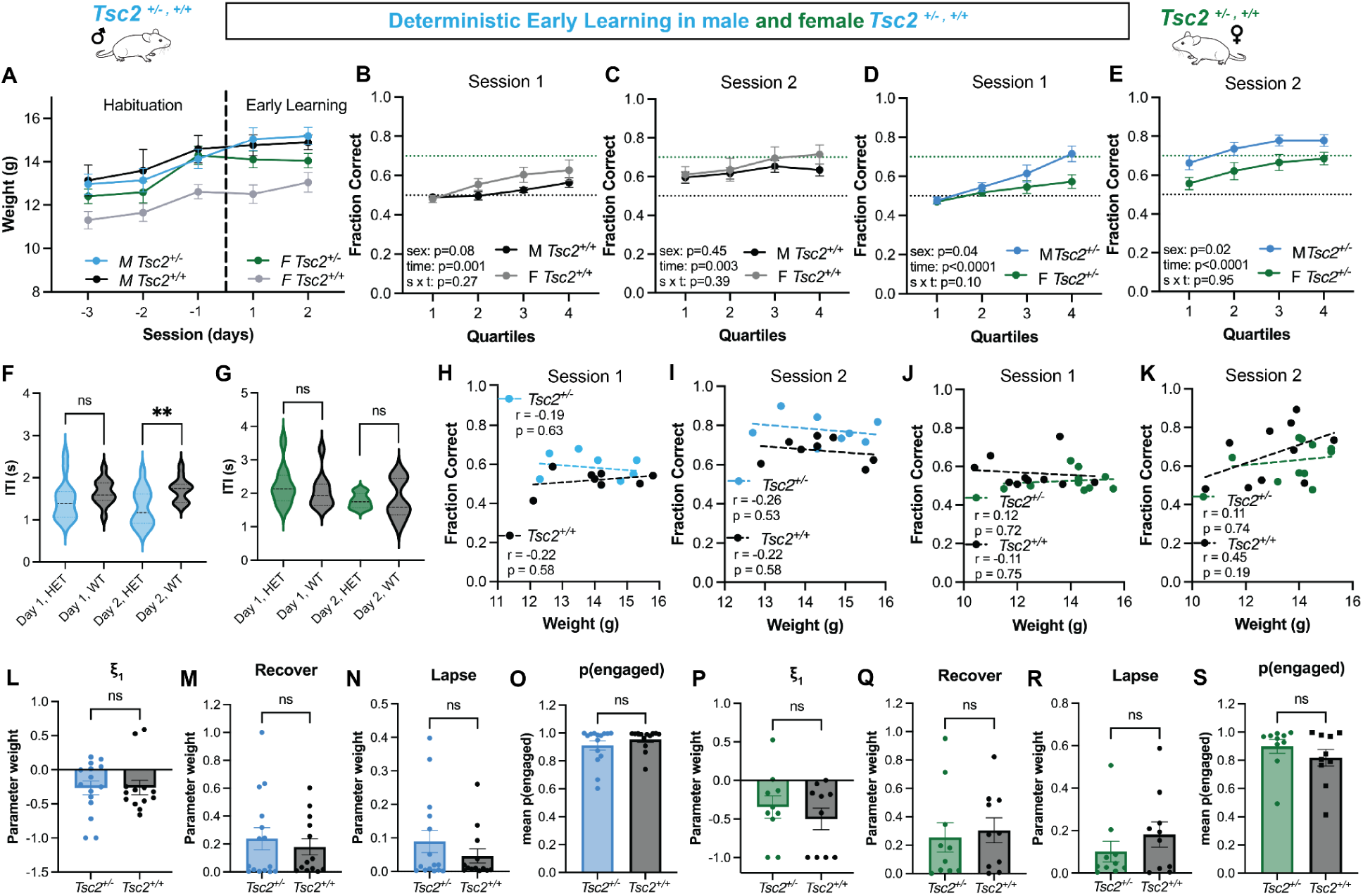
Additional analyses for *Tsc2* mice trained in the 2AFC with a deterministic schedule show sex differences in learning in Het, but not WT mice Additional modeling parameters to support the main text (Fig. 1 and Fig. 2) are also shown. (A) There was a significant gain in weight across habituation and early learning for all groups of animals (Male Het, n = 15; Male WT, n = 13; Female Het, n = 10; Female WT, n = 10). Mixed-effect model, using REML, fixed effect of time: *F*(2.59, 102.3) = 36.50, *p* < 0.0001; fixed effect of group: *F*(3, 40) = 4.30, *p* = 0.01. (B-C) Comparison of male (black) and female (grey) *Tsc2* WT animals in Session 1 and in Session 2 revealed no differences in performance or interaction. (D-E) However, there was a significant main effect of sex between male and female *Tsc2* Het for both Session 1 and Session 2. (F-G) Group average ITI comparisons for Session 1 and Session 2 for male *Tsc2* Hets (blue) and female *Tsc2* Hets (green) compared to sex-matched WTs (black) with pairwise comparison results described in the text. Note: n = 8 for WT females in Session 2 due to some missing ITI data. Violin plots show quartiles (25% and 75%), mean, and range. (H-I) Performance in Session 1 and 2 for male *Tsc2* Het or WT was not correlated with weight, using Pearson’s or Spearman’s correlation dependent on normality tests. (J-K) Neither *Tsc2* WT or Het female mice had performance in the task that was correlated with weight, using Pearson’s or Spearman’s correlation dependent on normality tests. (L-N) Male *Tsc2* Het and WT groups showed no significant differences in additional model parameters, ξ_1_, recover, and lapse, all fit from the winning model. Others are reported in main figures and text. (O) In a post-hoc analysis, male *Tsc2* Het and WT animals showed a similar level of mean engagement throughout the task. (P-R) Female *Tsc2* Het and WT mice showed no significant group differences in additional model parameters (ξ_1_, recover, and lapse) fit from the winning model. Others are reported in main figures and text. (S) In a post-hoc analysis, female *Tsc2* Het and WT animals showed a similar level of mean engagement throughout the task. See results section of main text for full statistics. Error bars represent SEM.

**Supplemental Figure 2:**
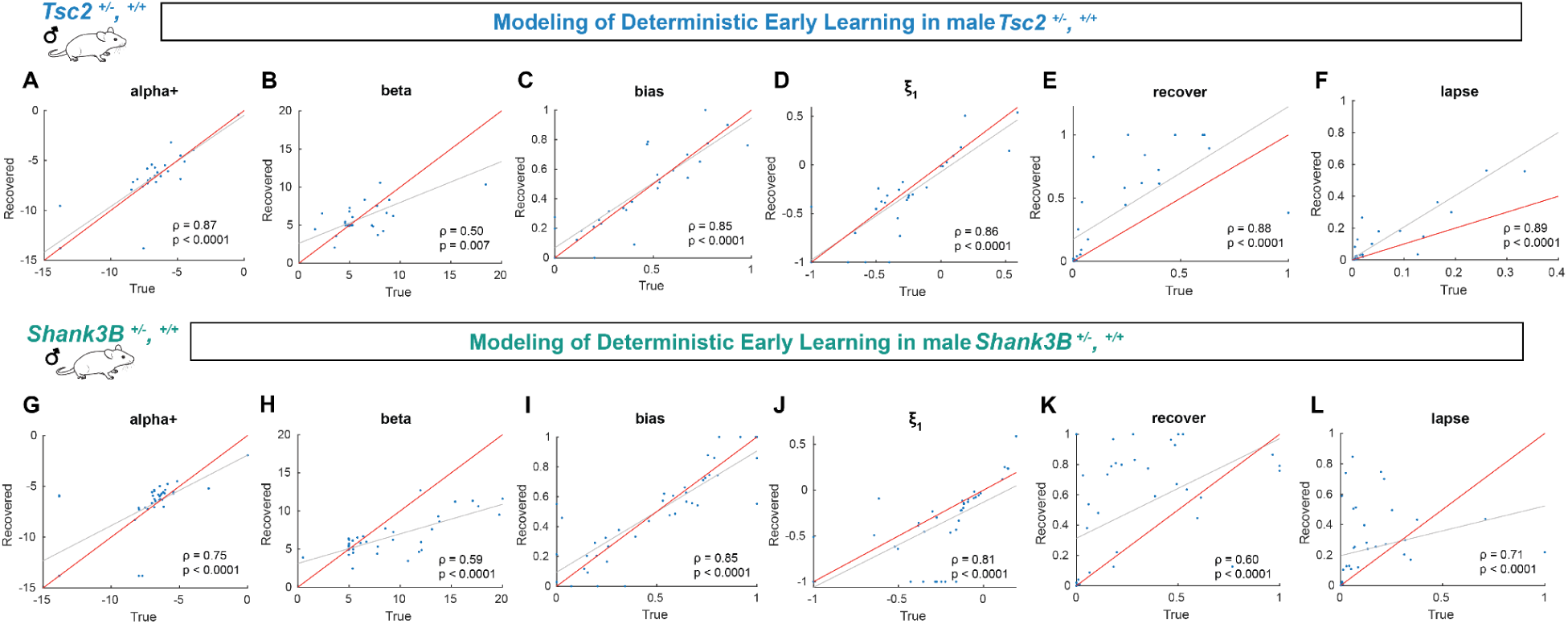
Parameters from winning model, a0bs1 hybrid, are identifiable. We tested a series of models that explored which strategies mice may use to learn the odor-based 2AFC task in Session 1. The winning model (as determined by BIC, see Figure 2 and Supplemental Figures 4 and 5) had 6 free parameters that included an *α*+ (learning from positive outcomes), an inverse temperature parameter □, a bias parameter that described an animal’s likelihood to continuously choose one side, a stickiness parameter, ξ_1_, that indicated the animal’s likelihood to repeat the previous trial’s action, and two parameters, lapse and recover, that marked the transition probabilities between an engaged and a disengaged (biased) state. In order to validate the winning model we generated data by simulating the model with parameters fit to individual sessions and fit the simulated data to obtain recovered parameters. We found that generated parameters (x-axis) were highly correlated with recovered parameters (y-axis) for both *Tsc2* (A-F) and *Shank3B* (G-L) male Session 1 data. This correlation indicates that the model parameters are identifiable (Wilson & Collins, 2019).

**Supplemental Figure 3:**
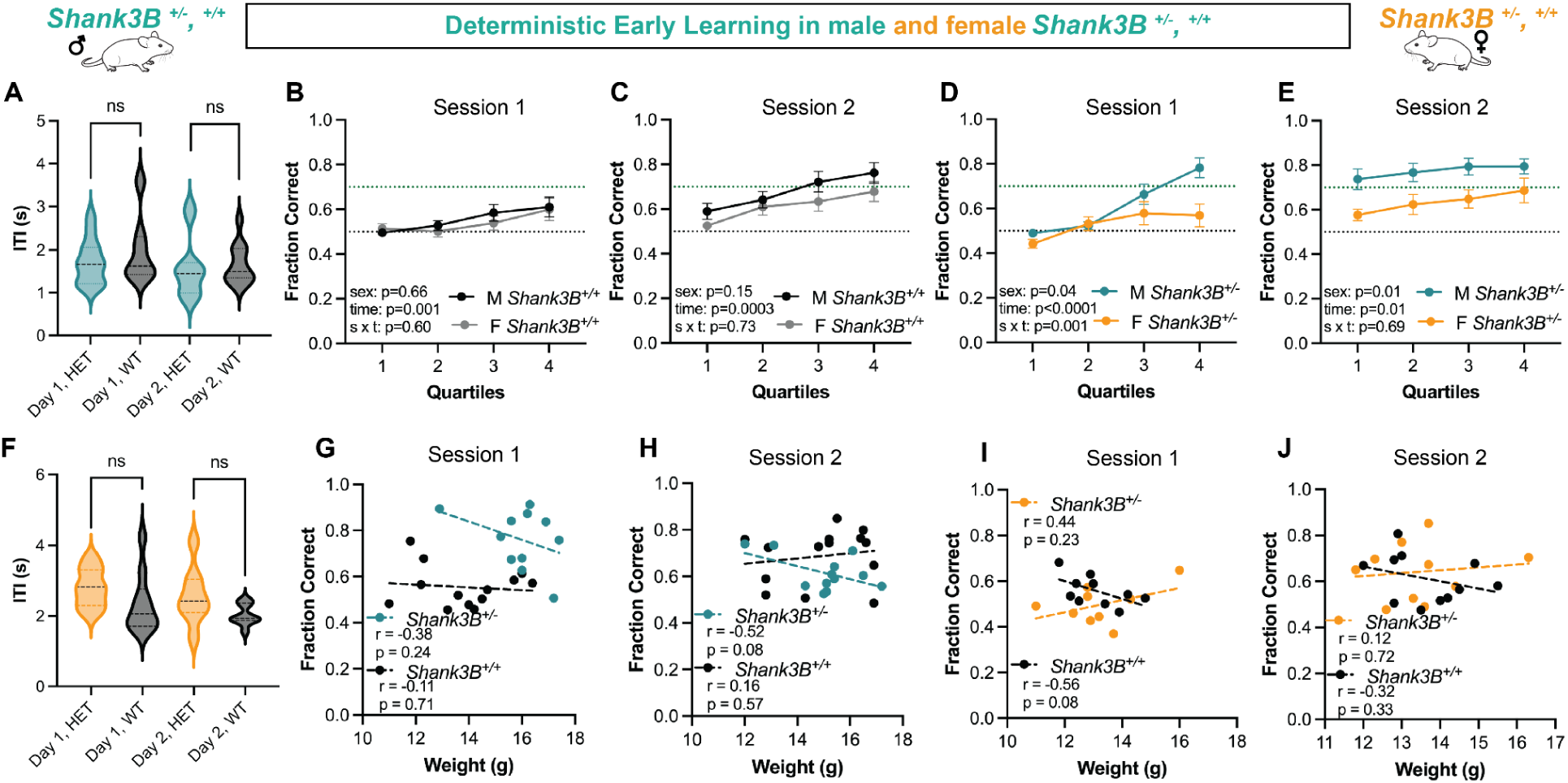
Additional analyses of male and female *Shank3B* mice trained in the 2AFC with a deterministic schedule show sex differences in learning in Het, but not WT mice. (A) Average ITI was comparable between Het (teal) and WT (black) in Session 1 (Het, n = 12; WT, n = 14) and Session 2 (Het, n = 11; WT, n = 13). (B) Comparison of male (n = 14) and female (n =11) *Shank3B* WT animals in Session 1 and in Session 2 (C) revealed no sex differences in learning performance. However, when comparing male (n = 12) and female (n = 11) *Shank3B* Hets, there was a significant main effect of sex in Session 1(D) and Session 2 (E) as well as a significant sex and time interaction between male and female Hets in Session 1. (F) Average ITI was comparable for female Shank3B Het (yellow) and WT (black) mice in Session 1 (Het, n = 9; WT, n = 10) and Session 2 (Het, n = 9; WT, n = 9). There were no significant correlations between fraction correct performance and weight in Session 1 and Session 2 for males (G-H) or for females (I-J).

**Supplemental Figure 4:**
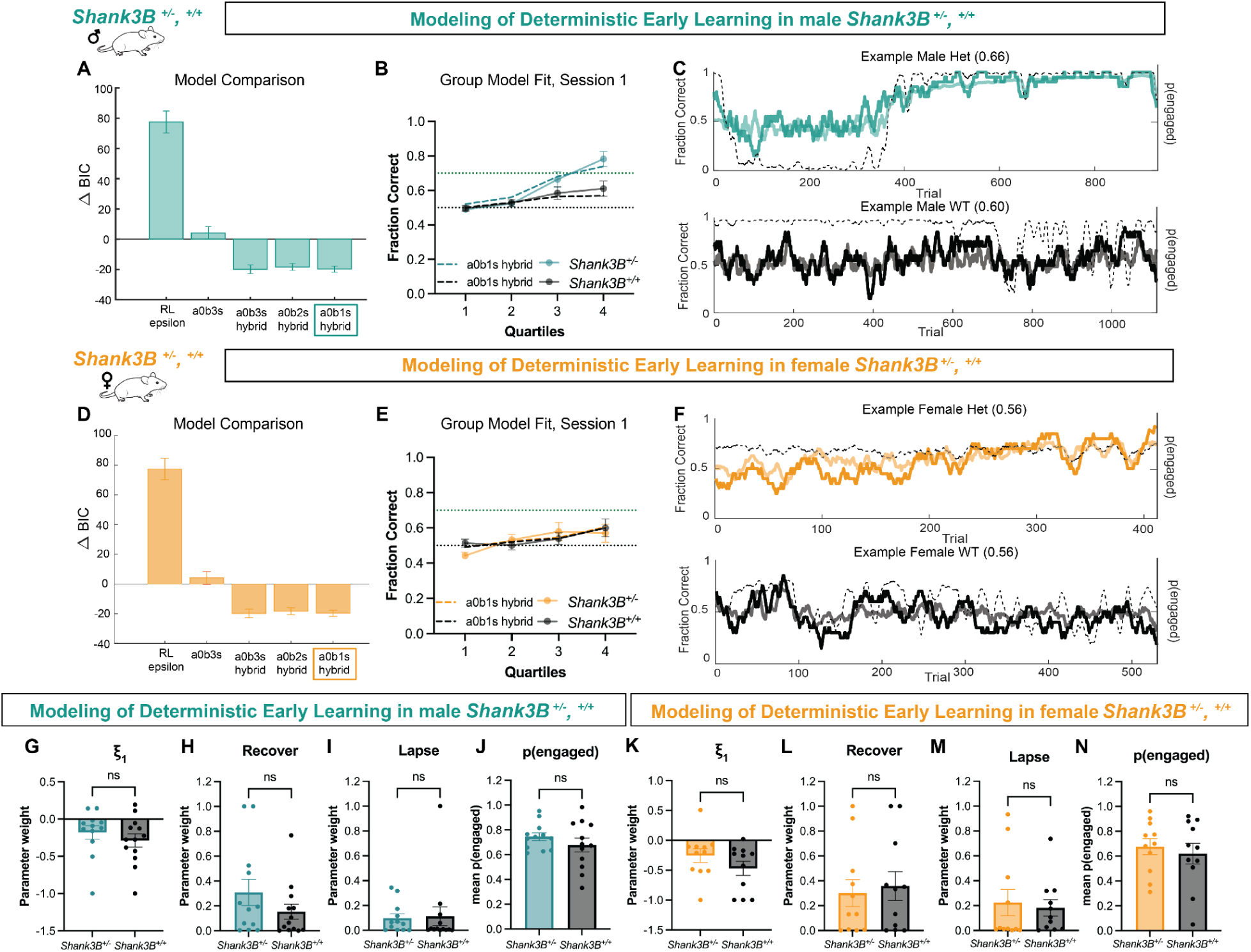
Additional data from male and female *Shank3B* Het and WT mice trained with a deterministic schedule in the 2AFC task: model comparison, validation, and additional fit parameters. (A,D) We compared a series of reinforcement learning and mixture models that can account for periods of task disengagement at the policy level. Using Bayesian Information Criterion (BIC), we determined that models that contained the mixture policy (hybrid) had a lower BIC score. The change in BIC is calculated by the difference in each model’s BIC and the mean BIC of all models examined. All hybrid models were comparable, but the model with the lowest BIC (colored square) was a0b1s_hybrid for both males (A) and female (D) Shank3B. (B,E) We generated data by simulating the model with parameters fit on individual sessions. Then, we inspected the group similarity between simulated data (teal dotted line for male *Shank3B* Het and yellow dotted line for female *Shank3B* Het and black dotted line for WTs) and found that the model simulated with fit model parameters captures group mouse learning curve data. (C,F) Examples of individual trial-by-trial model fits for both males (C) and females (F) with the teal or yellow line reflecting male or female trial-by-trial performance, respectively, teal or yellow transparent line reflecting the model fit, and the thin black dashed line reflecting the likelihood that the animal is in an engaged state. (G-I) Male *Shank3B* Het and WT mice showed no significant differences in additional model parameters (ξ_1_, recover, and lapse) fit from the winning model, pairwise comparison results described in main text results. (J) In a post-hoc analysis, male *Shank3B* Het and WT animals showed a similar level of mean engagement throughout the task. (K-M) Female *Shank3B* Het and WT mice showed no significant differences in additional model parameters (ξ_1_, recover, and lapse) fit from the winning model, pairwise comparison results described in main text results. (N) In a post-hoc analysis, female *Shank3B* Het and WT animals showed a comparable level of mean engagement throughout the task. Error bars throughout represent SEM.

**Supplemental Figure 5:**
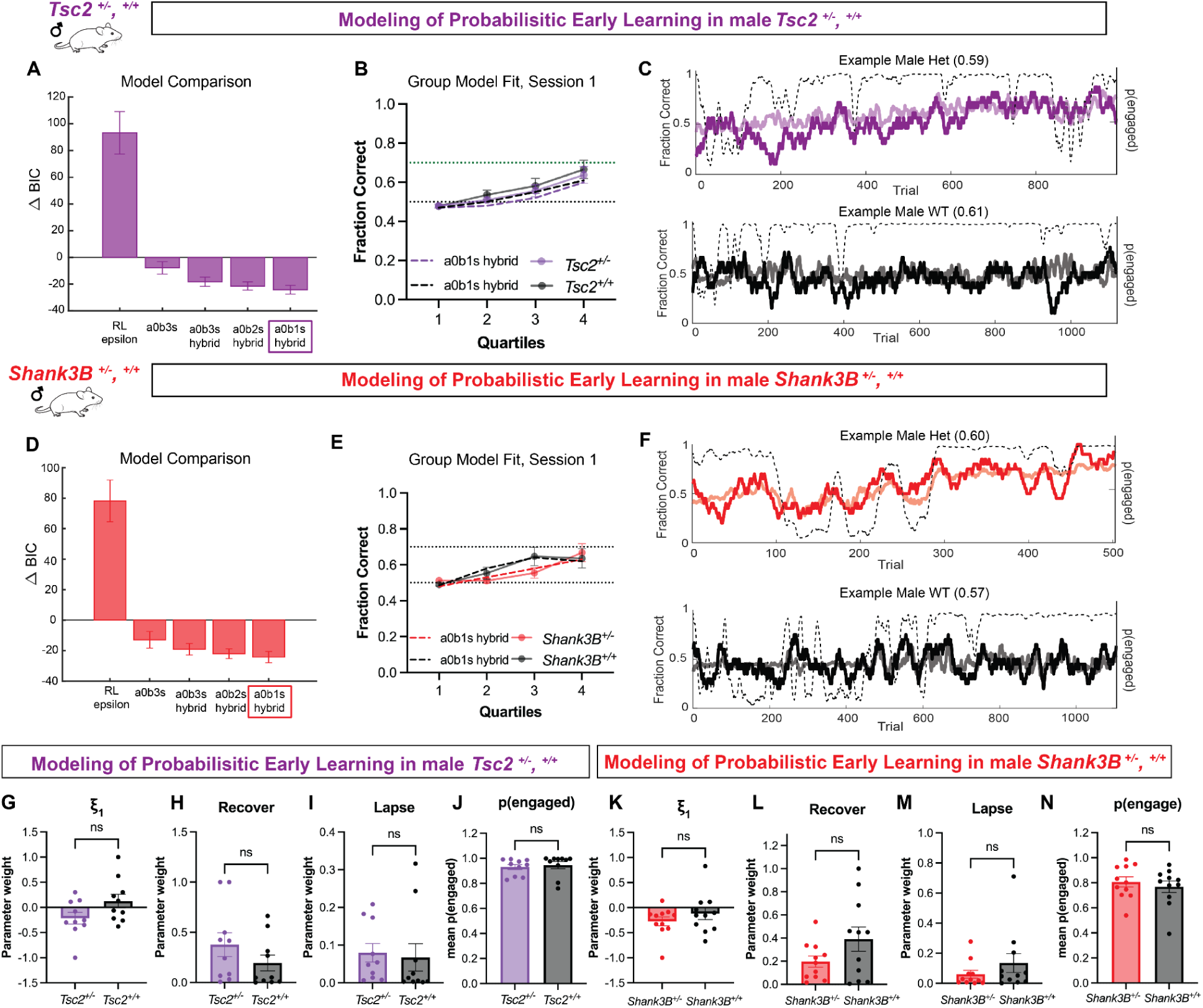
Additional data from male *Tsc2* and *Shank3B* Het and WT mice trained in the 2AFC task with a probabilistic schedule: model comparison, validation, and additional fit parameters. (A,D) We examined a series of mixture models as described in Supplemental Figure 4 and found the winning model to be a0b1s_hybrid for both male *Tsc2* (A) and male *Shank3B* (D) mice tested in the probabilistic schedule (where 10-20% of correct trials were not rewarded). This was the same winning model found above for the deterministic schedule (when 100% of correct trials rewarded) (B,E) We generated data by simulating the model with parameters fit on individual sessions. Then, we inspected the group similarity between simulated data (purple dotted line for male *Tsc2* Hets and red dotted line for male *Shank3B* Hets) and found that the model simulated with fit model parameters captures group mouse learning curve data. (C,F) Examples of individual trial-by-trial model fits for both *Tsc2* Het males (C) and *Shank3B* Het males (F) with the purple or red lines reflecting male *Tsc2* or *Shank3B* trial-by-trial performance, respectively, the light/transparent purple and red lines reflects the model fit and the thin black dashed line is the likelihood that the animal is in an engaged state. (G-I) Male *Tsc2* Het and WT mice had no differences in additional model parameters (ξ_1_, recover, and lapse) fit from the winning model, pairwise comparison results described in text. (J) In a post-hoc analysis, male *Tsc2* Het and WT animals showed a similar level of mean engagement throughout the task. (K-M) Male *Shank3B* Het and WT mice had no differences in additional model parameters (ξ_1_, recover, and lapse) fit from the winning model, pairwise comparison results described in main text. (N) In a post-hoc analysis, male *Shank3B* Het and WT animals showed a similar level of mean engagement throughout the task. Full statistics reported in main text and error bars represent SEM.

**Supplemental Figure 6:**
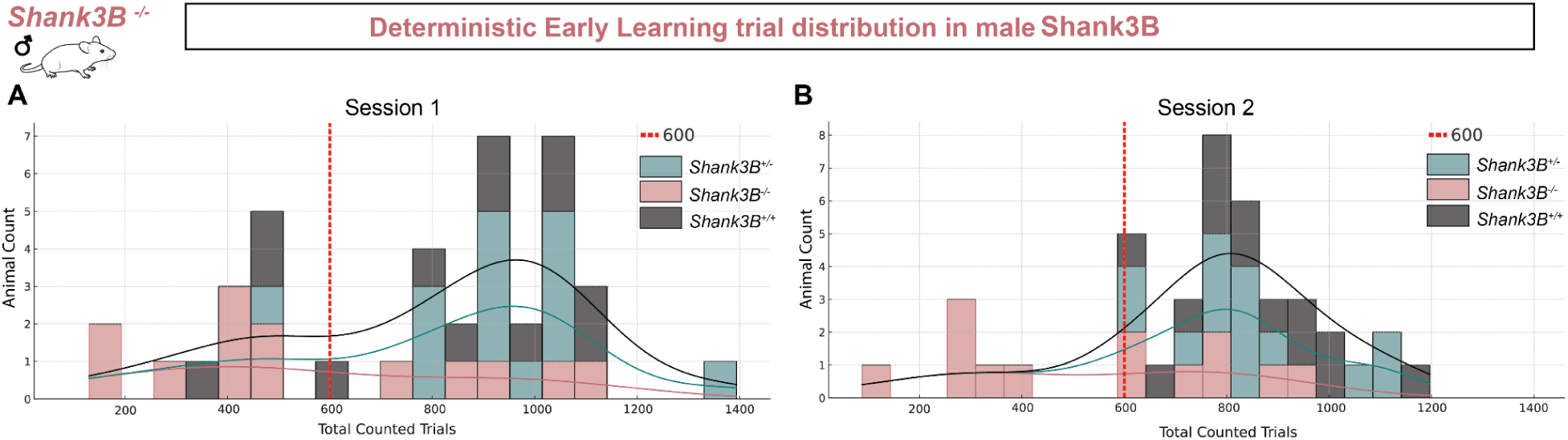
Distribution of trial counts for deterministic early learning in male Shank3B mice. (A-B) Histograms showing the total counted trials for male *Shank3B* WT (black), Het (teal), and KO (salmon) mice across (A) Session 1 and (B) Session 2 of the deterministic task. Data are binned by 50 trials. Overlaid density curves represent the distribution within each group. The dashed red line indicates the 600-trial threshold. Differences in trial distribution highlight variation in trials completed among genotypes.

**Supplemental Figure 7:**
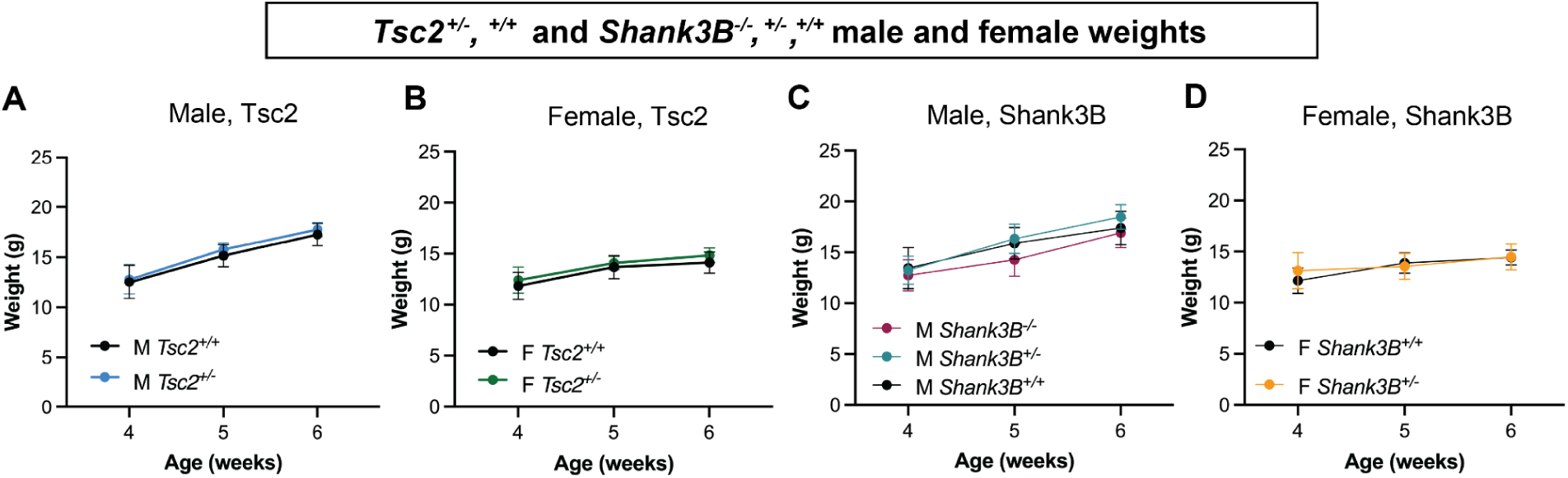
Growth trajectories in male and female *Tsc2* and *Shank3B* mice by genotype. We examined whether there was any difference between WT and Het weights for *Tsc2* and *Shank3B* mice across weeks 4, 5, and 6. Note that mice were tested in our task between weeks 4 and 5. Statistical comparisons were made using a 2-way ANOVA with factors for group (genotype) and week (4, 5, and 6), including the interaction between group and week. (A) There was no significant group difference between *Tsc2* WT and Het males (main effect of group: *F*(1, 58) = 2.37, *p* = 0.12; interaction: *F*(2, 58) = 0.10, *p* = 90), but there was a main effect of time (*F*(2, 58) = 90.3, *p* < 0.0001) indicating weight gain over time. (B) For female *Tsc2* WT and Hets, there was also no significant group difference or interaction (main effect of group: *F*(1, 50) = 3.92, *p* = 0.05; interaction: *F*(2, 50) = 0.11, *p* = 0.89). However, there was a significant main effect of time (*F*(2, 50) = 23.33, *p* < 0.0001), reflecting weight gain across weeks. (C) There was a significant main effect of genotype for male *Shank3B* mice (*F*(2, 105) = 7.78, *p* = 0.0007). Post-hoc Tukey tests showed that KO mice weighed significantly less than Het mice at Weeks 5 and 6 (*p* < 0.05) and less than WT mice at Week 5 (*p* < 0.05). No significant differences were observed between WT and Het at any time point. A significant main effect of time (*F*(2, 105) = 78.10, *p* < 0.0001) indicated weight increases across weeks, consistent across genotypes (no significant interaction: *F*(4, 105) = 1.23, *p* = 0.30). (D) For female WT and Het *Shank3B* mice there was significant weight gain across weeks (main effect of time: *F*(2, 60) = 11.47, *p* < 0.0001) but there was no significant difference in weight or weight gain between genotypes (main effect of genotype: *F*(1, 60) = 0.55, *p* = 0.46; genotype and time interaction: *F*(2, 60) = 1.60, *p* = 0.20). Data represents mean ± SEM.

**Supplemental Figure 8:**
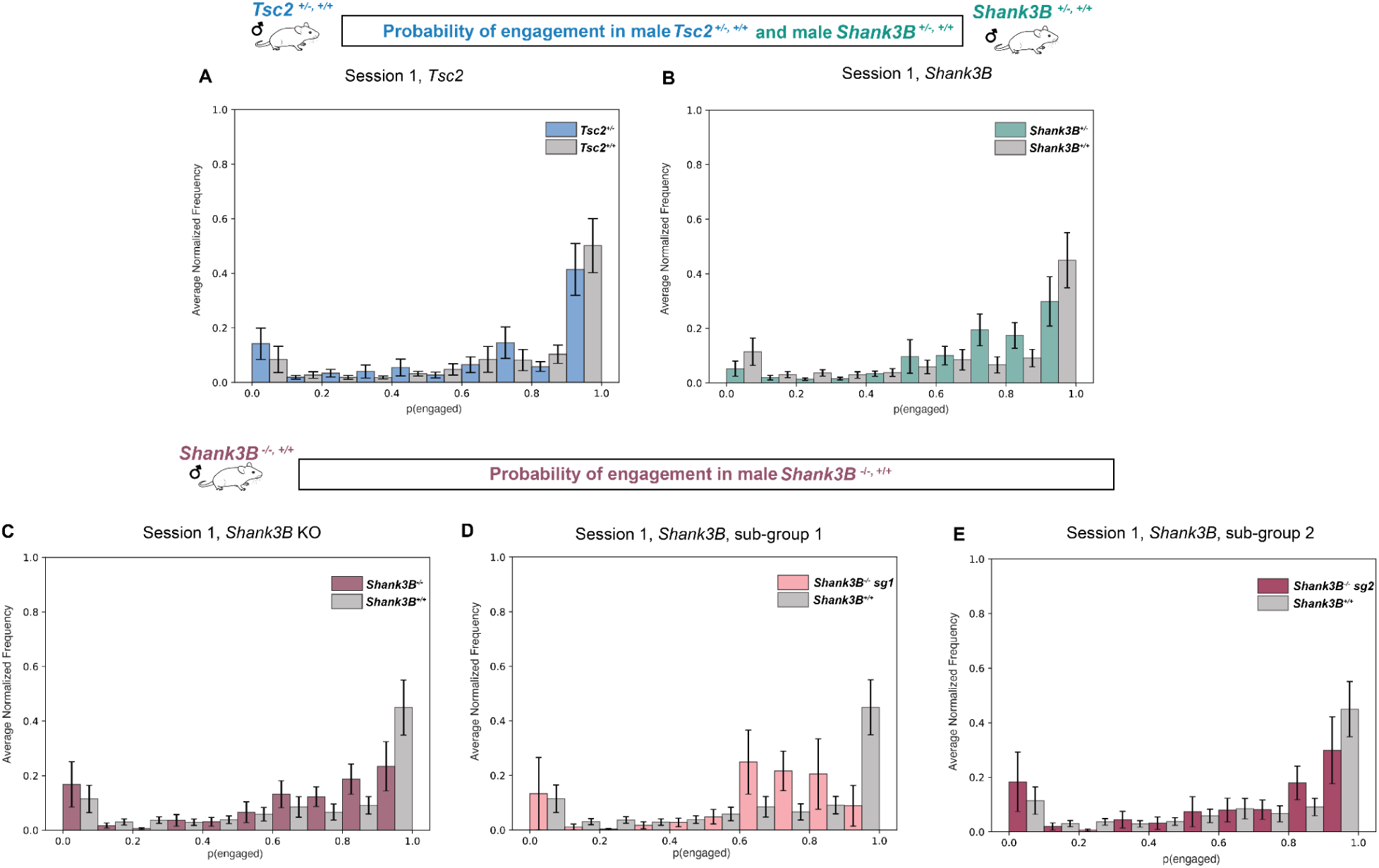
In Session 1 of the deterministic version of the 2AFC task, male *Tsc2* Het, *Shank3B* Het, and *Shank3B* KO mice showed similar distributions of engagement probability when compared to WT counterparts. To quantitatively assess differences in group distributions of p(engaged), we computed the Kullback-Leibler (KL) divergence for each pair of group-level distributions and performed a permutation test (see Materials & Methods). KL divergence measures the difference between two probability distributions, with lower values indicating higher similarity and values close to zero suggesting negligible divergence. Each panel shows the average normalized frequency that each group spent in p(engaged) bins during the first session. (A) *Tsc2* WT and Het males showed a low observed KL divergence of 0.129, with a non-significant *p*-value of 0.433. (B) *Shank3B* WT and Het males had an observed KL divergence of 0.264, p = 0.20, further indicating high overlap in their distributions. We also analyzed *Shank3B* KO males as a single group and divided into subgroups described in the text, compared to *Shank3B* WT males. (C) The comparison between *Shank3B* WT and all *Shank3B* KO males yielded a KL divergence of 0.241, *p* = 0.266. (D) *Shank3B* WT and *Shank3B* KO subgroup 1 had an observed KL divergence of 0.601, *p* = 0.161, while (E) *Shank3B* WT and *Shank3B* KO subgroup 2 had an observed KL divergence of 0.121, *p* = 0.766. Together, these results highlight that the KL divergence values remain low across comparisons and, coupled with non-significant *p*-values, suggest no meaningful differences in the group-level distributions of p(engaged). Error bars represent SEM.

